# Tunable hetero-assembly of a plant pseudoenzyme-enzyme complex

**DOI:** 10.1101/717082

**Authors:** Irina V. Novikova, Mowei Zhou, Chen Du, Marcelina Parra, Doo Nam Kim, Zachary L. VanAernum, Jared B. Shaw, Hanjo Hellmann, Vicki H. Wysocki, James E. Evans

## Abstract

Pseudoenzymes have emerged as key regulatory elements in all kingdoms of life despite being catalytically non-active. Yet many factors defining why one protein is active while its homolog is inactive remain uncertain. For pseudoenzyme-enzyme pairs, the similarity of both subunits can often hinder conventional characterization approaches. In plants, a pseudoenzyme PDX1.2 positively regulates vitamin B_6_ production by association with its active catalytic homologs such as PDX1.3 through an unknown mechanism. Here we used an integrative experimental approach to learn that such pseudoenzyme-enzyme pair associations result in hetero-complexes of variable stoichiometry, which are unexpectedly tunable. We also present the atomic structure of the PDX1.2 pseudoenzyme as well as the population averaged PDX1.2-PDX1.3 pseudoenzyme-enzyme pair. Finally, we dissected hetero-dodecamers of each stoichiometry to understand the arrangement of monomers in the hetero-complexes and identified symmetry-imposed preferences in PDX1.2-PDX1.3 interactions. Our results provide a new model of pseudoenzyme-enzyme interactions and their native heterogeneity.

## Introduction

Pseudoenzymes are homologs of enzymes that are catalytically deficient or inactive. Prior research from select protein families of kinases and phosphatases has shown that many pseudoenzymes retain the structural fold of their catalytically active partners and function as protein interaction modules^1^. Structural studies were the key tools in exploring and clarifying the molecular mechanisms of such systems and they uncovered their roles as allosteric regulators, scaffolds, molecular switches and substrate competitors^2-4^. These pseudoenzymes have to be diligently targeted, or anti-targeted (i.e. avoid intervention), for desirable outcomes in drug design or bioengineering applications so as to not be indiscrimately treated as their active homologs. To understand pseudoenzyme roles on a wider scale, genomic studies now show that pseudoenzymes account for 5-10% of the proteome and are associated in specialized signaling networks across all life domains^5^. However, the high structural similarities between pseudoenzymes and canonical enzymes create new experimental challenges in classical molecular and biochemical characterization approaches for unraveling the underlying mechanism. This is the case for the present study, which is focused around pseudoenzyme PDX1.2, a regulator of vitamin B_6_ biosynthesis in plants.

Vitamin B_6_ functions as a cofactor for over a hundred enzymatic processes and also provides an important defense mechanism against oxidative stress and pathogen infection^6-10^. Plants are able to synthesize vitamin B_6_ (PLP) *de novo* by two enzymes – PDX1 and PDX2 (Pyridoxine Biosynthesis 1 and 2)^6,11^. In particular, in *Arabidopsis thaliana*, there are three homologs of PDX1, designated as PDX1.1, PDX1.2 and PDX1.3, and one homolog of PDX2^11^. Of these, PDX1.1, PDX1.3 and PDX2 are all catalytically active enzymes while PDX1.2 has been shown to be an inactive variant^11^. *In vivo* experiments showed that PDX1.2 expression is upregulated by heat and oxidative stress and coincides with a boost in vitamin B_6_ synthesis, suggesting a positive regulatory impact^12-14^. PDX1.2 was first proven to interact with its catalytic PDX1.1 and PDX1.3 proteins *in vivo* and to form complexes of high molecular weight similar to its catalytic homologs^15^. Recombinant co-expression of PDX1.2 with either PDX1.1 or PDX1.3 showed dodecameric hetero-complexes^13^.

PDX1.2 protein resists crystallization protocols while PDX1.3 homo-complexes and PDX1.2/PDX1.3 hetero-complexes are compatible with macromolecular crystallography^16^. However, unlike PDX1.3 homo-complexes, the crystals of PDX1.2/PDX1.3 hetero-complexes sufferred from a statistical disorder, where the positions of individual proteins were not distinguished despite exhaustive approaches used^16^. The ambiguity in the data prohibited the determination of the assembly mechanism but an interhexamer assembly mechanism was proposed as the likely cause of interaction where one hexameric ring of active PDX1.1 (or PDX1.3) stacks on top of a hexamer ring of inactive PDX1.2 (Fig.1a) to create a final hetero-dodecameric state^13,16^. Herein, using an integrated biochemical and structural approach combining cell-free protein synthesis technology, native mass spectrometry (native MS), and cryo-electron microscopy (cryo-EM), we show that PDX1.2 and PDX1.3 form hetero-dodecamers through intrahexamer association of two symmetric or similar hexamers with varying but tunable stoichometry.

**Fig. 1.**
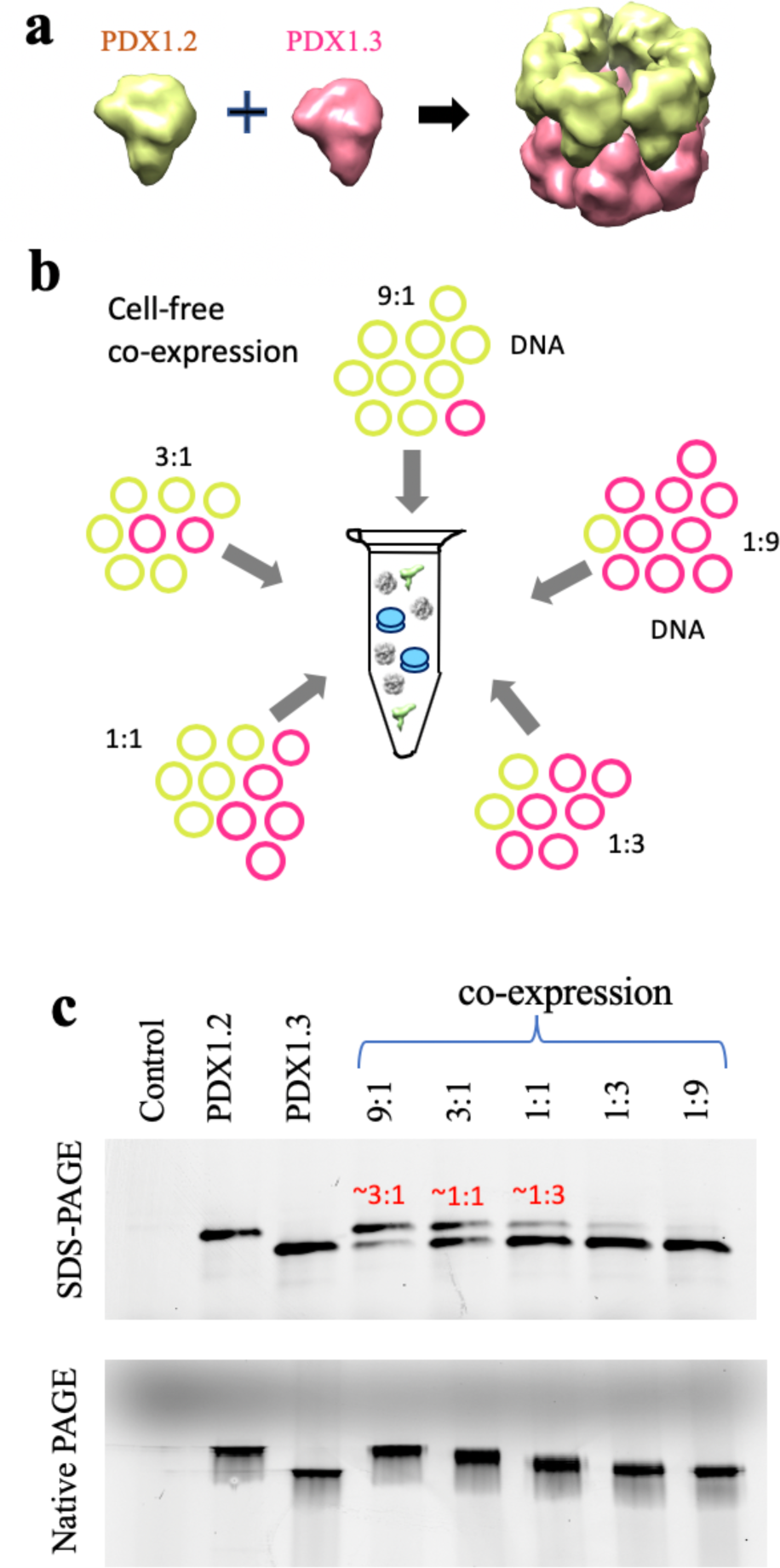
PDX1.2 and PDX1.3 co-expression in a test tube. **a**. Schematic showing the prior proposed hetero-assembly of PDX1.2-PDX1.3 association. **b.** Illustration of experimental design where variable amounts of DNA plasmids are fed into the cell-free protein translation system to control and study the co-expression complexes. **c**. PAGE analysis of newly-synthetized PDX proteins, detected by fluorescence in a crude mixture. The control sample is a translation reaction where DNA plasmid is not supplemented. PAGE data include a denaturing SDS-PAGE gel on the top and Native PAGE on the bottom. In the co-expressed samples, protein molar ratios, determined by in- gel quantifications, are shown in red. Co-expressed conditions are denoted as 9:1, 3:1, 1:1, 1:3 and 1:9, where these numbers correspond to DNA template ratios of PDX1.2 to PDX1.3.

## Results

### Cell-free expression of PDX homo/hetero-complexes

We employed a cell-free protein expression pipeline using wheat germ extract^17-19^ to provide the closest translational environment for the production of these plant proteins in terms of folding and post-translational modifications. The “open-format” of a cell-free platform provided us with an opportunity for precise stoichiometric control of protein co-expression (Fig.1b) by varying the amounts of corresponding DNA templates, not easily achieved by cell-based protein expression methods. We prepared two DNA plasmids constructs (one for PDX1.2 and one for PDX1.3), and combined them in specific molar ratios to achieve the desired protein output in the co-expression conditions (Fig.S1).

In the initial cell-free translation experiments, we supplemented all the translation reactions with fluorophore-labeled lysine-charged tRNA as recently described^19^. This procedure allowed detection of newly-synthesized proteins in the crude mixture with no need for purification. For co-expression, different DNA template ratios (9:1, 3:1, 1:1, 1:3 and 1:9) were tested to establish that the molar ratio of expressed proteins can be controlled in a precise fashion. For instance, by in-gel quantification, the co-expression conditions 9:1, 3:1 and 1:1 correspond to protein product compositions of 3:1, 1:1 and 1:3 (Fig.1c). The same samples were also analyzed under native gel conditions where all co-expressed samples traveled faster than individual PDX1.2 homo-complexes but slower than PDX1.3 homo-complexes. In addition, the bands for co-expression complexes appeared significantly wider in size suggesting the presence of multiple stoichiometric dodecamer species. The mobilities of these samples also appeared to be dependent on the protein composition where a higher proportion of PDX1.3 in the co-expressed sample caused faster migration of the hetero-complex, and the opposite was true for higher PDX1.2 content. The variable hetero-complex migration on the gel and the thickness of the bands do not support the previously proposed interhexamer assembly mechanism^16^ where a defined single band of a single stochiometric ratio of 6:6 would be expected. Instead, PDX1.2 and PDX1.3 likely form mixed hetero-complexes.

We also verified that co-expression was essential for the PDX1.2 and PDX1.3 hetero-complex formation, supporting previous discoveries^13,16^. Mixing the two separately expressed and purified proteins post-translationally did not produce detectable amounts of hetero-complexes on native PAGE (Fig. S2a). The same co-expression experiments were conducted using active PDX1.1 enzyme (instead of PDX1.3) and PDX1.2, and identical pseudoezyme-enzyme assembly behavior was observed (Fig.S2b). Because PDX1.3 is the dominant homolog in *A. thaliana*^20,21^ and was previously found to be more impacted enzymatically by the presence of PDX1.2^13^, we have limited our further structural survey to PDX1.2 and PDX1.3.

### Tunable stochiometry and activity of PDX hetero-complexes

For further structural characterization, we scaled-up the synthesis and then purified select samples: PDX1.2 and PDX1.3 homo-complexes and hetero-complexes at the co-expression conditions 9:1, 3:1 and 1:1 (Fig. S3). Protein sequences were verified by peptide mapping (Fig. S4). Liquid chromatography mass spectrometry (LCMS) of intact (denatured) proteins showed masses of 37339.7 and 36405.5 Da for PDX1.2 and PDX1.3, respectively (Fig. S5), which are consistent with their monoisotopic theoretical masses of 37339.6 Da and 36405.3 Da. Although approximately 50% of the protein N-termini were found to be acetylated, no preferential acetylation profile was detected suggesting these post-translational modifications are unlikely to play a role in PLP biosynthesis regulation (Fig. S5-S6).

While the native PAGE results indicated the presence of mixed co-complexes in each co-expression condition, the technique did not have the resolution to define the stoichiometry. We therefore employed native MS, where we electrosprayed the proteins under nondenaturing conditions in 200 mM ammonium acetate solution. The intact 12mers were detected around *m/z* of 9000, carrying ∼ 50 positive charges (Fig. S7). All species in the co-expressed samples were resolved and the stoichiometry could be accurately determined from the measured mass, as shown by the native MS spectrum of co-complex 9:1 in Fig. 2a. From the spectra, mass distributions of all species in the sample can be deconvoluted by combining the intensities across charge states (Fig. 2b). As expected, the PDX1.2 and PDX1.3 samples showed predominantly homo-12mers with molecular weights of 449 kDa and 437 kDa, respectively. Each co-expressed sample generated 6-8 hetero-complexes of variable stoichiometry that were all dodecameric and influenced by the initial plasmid DNA ratio input to the cell-free reaction. For example, the 9:1 co-expression condition resulted in a pool of hetero-complexes with PDX1.2 representing the major protein constituent and spanning observed stoichiometry values in the 6:6 to 12:0 range (6:6, 7:5, 8:4, 9:3,10:2, 11:1 and 12:0). For the 3:1 co-expression condition, there is a shift to the 3:9 - 11:1 stoichiometry range, while the 1:1 co-expression condition was limited to the 0:12 - 8:4 stoichiometry range (Fig. 2b). The observed stoichiometry values were consistent with the amounts of proteins produced in each sample (see Fig.1c and Fig. S3). By varying the input DNA ratio (and thus protein ratios), the full range of co-complexes from 0:12 to 12:0 stoichiometry can be sampled, demonstrating the tunability of these protein-protein associations.

**Fig.2.**
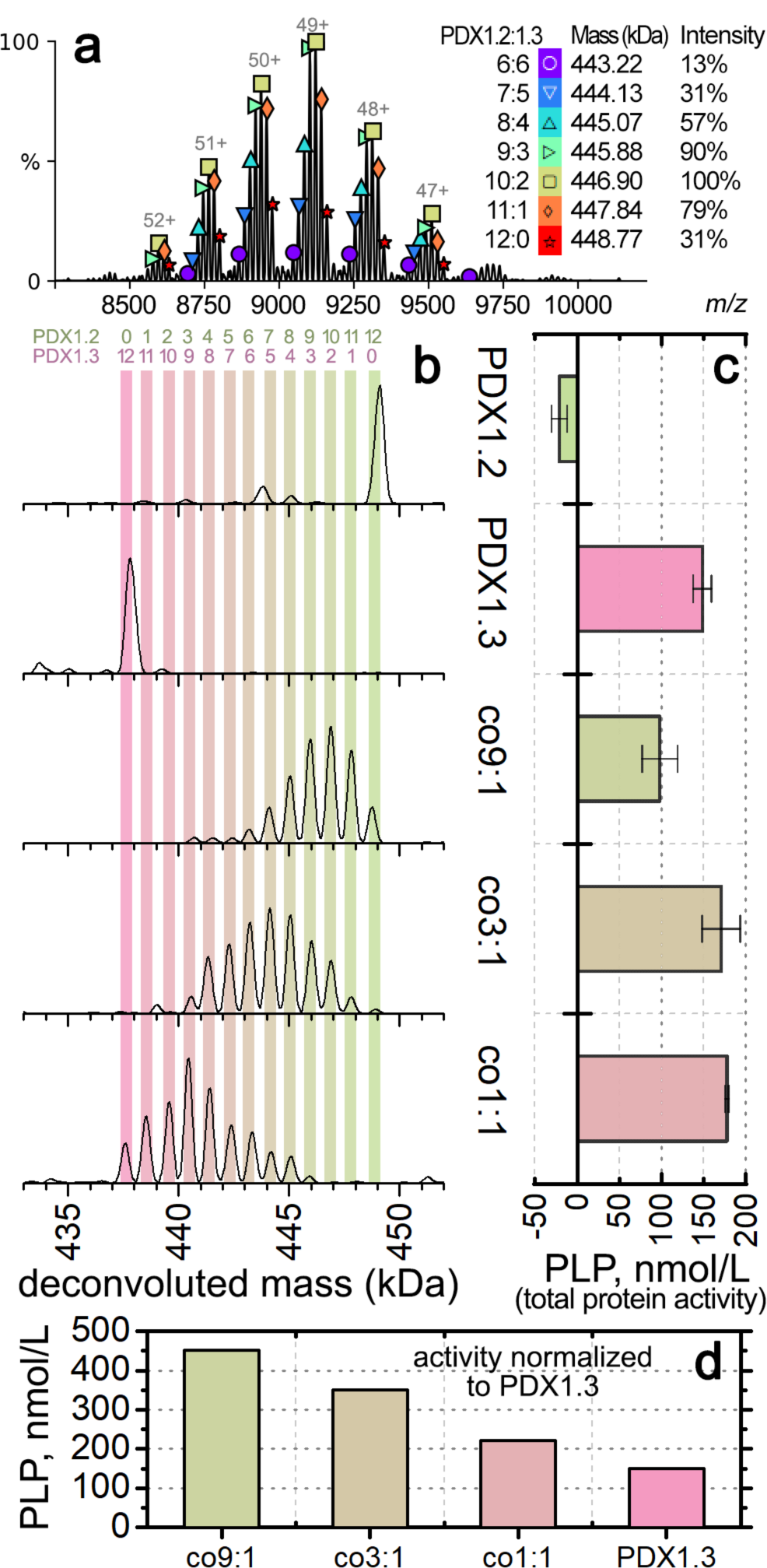
Oligomeric and enzymatic characterization of individual PDX homo- and hetero-complexes. **a**. Representative native MS spectrum for co-expression complex 9:1, zoomed into the 12mer region. Each symbol above the peak indicates one 12mer species, with their assignment, mass, and relative intensity shown on the right. Each peak with the same symbol is the same species carrying a different number of charges. Annotation was performed using UniDec. **b**. Deconvoluted mass distributions of PDX 12mer complexes from different samples as labeled on the right. Color bars are overlaid to show the numbers of PDX1.2 and PDX1.3 protomers in the PDX 12mers as noted by the numbers on the top. **c**. Histograms showing the enzymatic activities based on total PDX protein concentration. Error bars represent the standardard deviations from triplicate experiments. The sample names match to the mass distributions shown horizontally in (b). **d.** Histograms showing the enzymatic activities normalized to the amount of PDX1.3 in each sample.

Interestingly, a significant amount of free monomer and dimer species were observed for PDX1.3 containing complexes in the low *m/z* region < 6000 (Fig. S7). In the co-expressed samples, the relative abundance of PDX1.3 monomers and dimers was also found to be positively correlated with the ratio of PDX1.3 in the hetero-complexes. In contrast, PDX1.2 appeared to be more stable under the same experimental conditions. We confirmed that 200 mM ammonium acetate did not cause noticeable protein unfolding or disassembly in solution prior to MS (Fig. S8). Previous native MS studies have shown that positively charged proteins may interact with silanol groups on borosilicate glass surface at sub-micron sized emitter tips, resulting in supercharging and unfolding during electrospray^22^. In addition, supercharging typically makes complexes easier to dissociate *in vacuo*^23,24^. Therefore, the dissociation of PDX1.3 containing 12mers might be due to unique charging properties of PDX1.3 that are distinct from PDX1.2.

To understand how the PDX1.2:PDX1.3 ratio in hetero-assembly affects their enzymatic performance, we employed a diagnostic assay where PLP is used as a cofactor molecule for apo-enzyme chemical transformations, the products of which were detected by a colorimetric approach (Fig.2c). Under these assay conditions with saturating substrate amounts, the specific activity of active enzyme PDX1.3 was determined to be 793 µmoles/min/mg protein. As expected, pseudoenzyme PDX1.2 was found to be inactive and displays a slightly inhibitory effect. When proteins are co-expressed under conditions 3:1 and 1:1, the presence of PDX1.2 hetero-complex positively impacts PLP synthesis with an increase of up to 120% in relative enzymatic activity. However, at 9:1, where 75% of the total protein is PDX1.2, the hetero-complex displays a decline in relative activity to 67% of that observed for PDX1.3 homo-12mer. Yet, the measured activities represent a sum of PDX1.2 and PDX1.3. If these relative enzymatic activities are normalized based on the amount of catalytically active PDX1.3 in each co-expressed sample, the positive regulatory impact of PDX1.2 is observed for every condition (Fig. 2d). The higher ratio of PDX1.2 incorporation showed higher impact on PDX1.3 catalytic activity with the corresponding outputs: 3-fold increase for co-expression condition 9:1, 2.4-fold increase for 3:1 and 1.5-fold increase for 1:1 (Note that co-expression conditions 9:1, 3:1, and 1:1 correspond to protein ratios 3:1, 1:1, and 1:3, respectively).

### Lateral symmetry within the seemingly stochastic assembly

To resolve the structures of the purified complexes we employed single particle cryo-EM. The PDX1.2 sample was first vitrified on EM grids and imaged in super-resolution mode using a Titan Krios cryo-EM with K2 direct electron detector (Fig. S9). A homogenous distribution of particles in various orientations are clearly visible in the thin layer of ice, and several rounds of 2D classification produced well-defined 2D classes where a stacked 2-ring architecture was easily distinguished and consistent with the expected dodecameric fold known for catalytically active PDX1.1 and PDX1.3. The final 3D volume was reconstructed at 3.2Å resolution, and a PDX1.2 homology model was then docked and real space refined (Fig. S10, Table S1). Residues 29-288 were clearly resolved while N- (residues 1-28) and C-terminal (residues 289-313) regions appear rather flexible due to the lack of associated densities in the volume. We also obtained a high-resolution cryo-EM dataset for the 9:1 co-expression complex (Fig. S11). A well-defined two-ring fold seemed visually identical to the PDX1.2 homo-12mer. Various heterogenous, non-uniform and local refinements did not sort out the potential subclasses, which is not surprising due to variability of the stoichiometry of the co-expressed species, the similarity of PDX1.2 and PDX1.3 folds, and the small difference (<1kDa) between their monomer molecular weights. As a result of the lack of distinct subclasses, the 9:1 co-expression complex particle dataset was processed as a single 3D class, and non-uniform refinement yielded the final map. Both the PDX1.2 and PDX1.3 homomers can be fitted independently with high validation scores (Fig. S12, Table S1) to any of the monomer positions - similar to the result reported recently for X-ray structure^14^. In other words, the two different subunits cannot be easily distinguished in the hetero-complex, suggesting a stochastic hetero-assembly.

To better understand the distribution of the subunits within the hetero-complexes, we mass isolated individual heteromers of each stoichiometry and subjected them to surface induced dissociation (SID) in a modified mass spectrometer^25^. The charge states of the proteins were reduced by adding triethylammonium acetate in solution, which suppressed unfolding upon activation and allowed more informative dissociation products to be obtained in SID^26^ (Fig. S13). A representative SID spectrum for the 6:6 hetero-complex is shown in Fig. 3a. Collision of the precursor ions into the surface under controlled laboratory frame collision energy (5.6 keV) in SID caused a rapid increase in an internal energy for the 12mer, leading to the dissociation into subcomplexes. Our previous studies have shown that subcomplexes generated in SID are often from dissection of the weakest interfaces and are reflective of the native quaternary structure of protein complexes^27,28^. In contrast, collision-induced dissociation (commercially available as higher-energy collisional dissociation, HCD, in the instrument used here) results in activation of the 12mer via multiple low-energy collisions and, at most, only resulted in stripping of unfolded monomers at the maximum collision energy (10.5 keV) (Fig. S14). Among the released species in SID, the most abundant was the 6mer (Fig. 3b). The other species such as 4mers and 8mers were also detected but at lower abundance. The major products, 6mers, are likely the result of horizontal cleavage along the interface of two 6mer rings (Fig. 3c). Another minor pathway of two vertical cleavages while maintaining the lateral interactions may have generated the 4mer+8mer pair and also contributed partially to the signal of 6mers. We suspect the 5mers in both spectra may partially originate from secondary dissociation of 6mers and are not from direct cleavage of the 12mers because of the low abundance of the complementary 7mers. Herein, we focus on the major pathway for structural elucidation of the hetero-assemblies.

**Fig.3.**
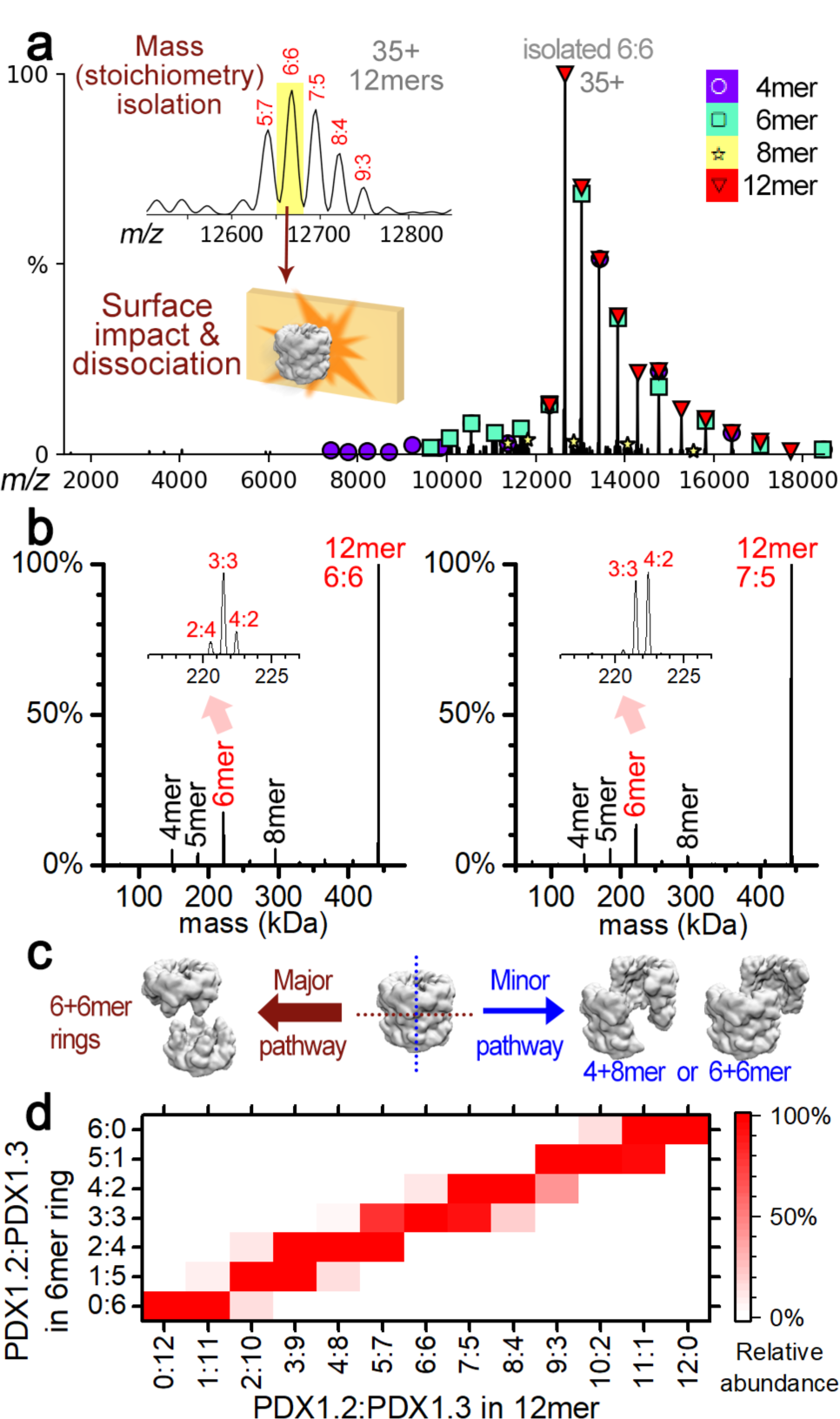
Dissection of PDX 12mers in the gas phase by SID. **a.** Representative SID spectrum for a heteromer with a single stoichiometry of 6:6 (PDX1.2:PDX1.3). Mass isolated 6:6 was collided with a surface to produce 6mers and other subcomplexes detected in the mass spectrum, which are labeled by the different symbols. The key to the symbols are on the top right. **b.** Deconvoluted mass distribution of the SID spectra for 6:6 (left) and 7:5 (right). Zoom-in views of the 6mer region are shown in the inserts. **c.** Proposed dissociation pathways for PDX by SID. The cleavage of the 12mer in the horizonal direction is a major pathway (left). A minor pathway for cleaving in the vertical direction likely produced several other species at low abundance. **d.** Heatmap showing the relative abundance (normalized to max) of the released 6mer rings (vertical axis) from each stoichiometry of 12mers (horizontal axis) following the major pathway. The contribution of 6mers from minor pathways were estimated by the abundance of 4/5/7/8mers, and was subtracted from the raw experimental data as described in the method section.

The deconvoluted SID spectra show a relatively monodispere stoichiometric distributions in the released 6mers. For example (Fig. 3b), ∼70% of the 6mers from the 6:6 heteromer had the stoichometry of 3:3 while the remainder were found to be 2:4 and 4:2. This suggests that the 6:6 12mers are formed in either the [3:3 and 3:3] or [2:4 and 4:2] combination of PDX1.2:PDX1.3. For 7:5 heteromer with odd numbers of each protein in the 12mer, 6mers with stochiometry of both 3:3 and 4:2 were released at similar intensities. The most abundant 6mers released by SID from other 12mers all have half of the stoichiometry of the 12mer precursors (Fig. S15). We estimated the abundances of the 6mers generated only from the major pathway (see methods for details) and plotted the relative abundances of the different stoichiometry as a function of the 12mer precursor stoichiometry in Fig. 3d. The narrow distribution of the stoichiometry of the 6mer rings strongly favors a lateral symmetry in the 12mers. If PDX1.2 and PDX1.3 were distributed stochastically, a wide variety of stochiometries should be observed in the released 6mers (Fig. S16).

### Similar protein folds but inverted electrostatic surface potentials

Because of the challenges in resolving individual proteins in co-complexes, we instead focused on the cryo-EM structure of the PDX1.2 homo-12mer for clues on its positive regulatory function by comparing it with several crystallographic structures of PDX1.3 available at different stages of enzymatic action^29^. Precursor binding stage PDX1.3-R5P (PDB:5lns), intermediate stage PDX1.3-I_320_ (PDB:5lnu), post-intermediate PDX1.3-I_320_-G3P (PDB:5lnw) and final product stage where PLP is still covalently bound to K166 (PDB:5lnr) were all aligned with our structure of PDX1.2 and overlaid for comparison (Fig. 4a). The PDX1.3 enzyme employs a lysine relay mechanism where all catalytic action happens within a single domain using two catalytic sites P1 and P2^29,30^. Two key residues K98 and K166 are at the heart of this process where they trap the substrates, covalently tether the intermediates and then shuttle the product out. We found that the overall monomer fold of PDX1.2 is very similar to PDX1.3, and the best fit for PDX1.2 is to the X-ray structure of PDX1.3-PLP, which has the P1 site unoccupied.

**Fig.4.**
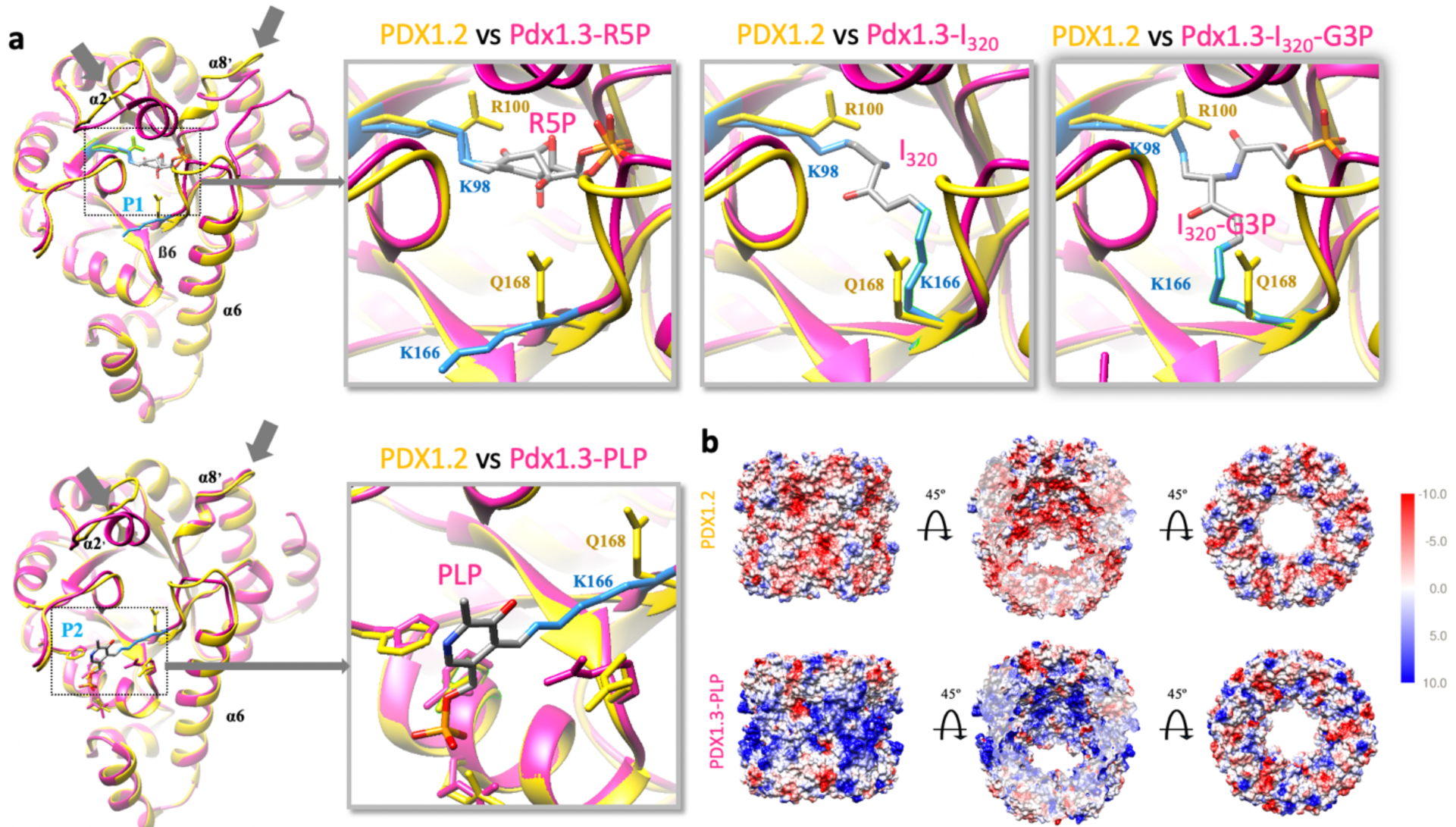
The cryo-EM structure of PDX1.2 versus the crystal structures of PDX1.3. **a**. Comparison of PDX1.2 individual subunit versus four different crystal structures, corresponding to the different stages of PLP synthesis at active sites P1 and P2. The PDX1.2 monomer subunit is in yellow, PDX1.3 subunit is in magenta. Key residues in PDX1.3 catalysis, K98 and K166, are shown in light blue. Corresponding residues in PDX1.2 include R100 and Q168 (in yellow). Substrate, intermediate products and PLP are colored by heteroatom. For structure-based sequence alignments and annotations, see Fig.S20. **b**. Coulombic electrostatic potential maps. The middle images highlight the sliced view to show the interior potential of the complex.

Importantly, the overall monomer fold of PDX1.2 is identical to PDX1.3 at the outer shell where proper higher order protein interactions with neighboring subunits need to be preserved (Fig. 4a). The conservation pressure on those amino acids, which are involved in quaternary interactions, appears not to be specific for *A. thaliana* and is imposed on many plant organisms (Fig. S17). Thus, the maintaince of PDX1.2 quaternary contacts with PDX1.3 is very important for pseudoenzyme functional performance. Major differences between PDX1.2 and PDX1.3 are observed in the area surrounding the catalytic site P1. Specifically, the **α2’** helix and **ß6-α6** loop are significantly altered in PDX1.2. The **α2’** helix, present in PDX1.3, is absent in PDX1.2 due to the amino acid truncation in that region and is substituted by a loop that points slightly outwards. The amino acid changes in the **ß6-α6** loop, including an amino acid insertion, create a slight kink in PDX1.2 architecture towards the center of the 12mer, resulting in the potential weakening/aborting of the phosphate binding at this site. Key lysine residues K98 and K166 which form covalent adducts with the intermediates of PLP formation in PDX1.3 are mutated to R100 and Q168 respectively in PDX1.2. While R100 is positioned similarly to K98 and can potentially form the imine bond with R5P as well, Q168 in PDX1.2 (in place of K166 in PDX1.3) points towards the P1 site similar to the intermediate state of the PLP synthesis instead of pointing to P2 as observed in the priming PDX1.3-R5P state. The same positional occupancy of Q168 was assigned to the composite assignment of Q168 and K166 in the crystallized PDX1.2-PDX1.3 co-complex where individual protein positions were not distinguished^16^. Such structural alterations in the P1 site in PDX1.2 are in agreement with the experimental data, showing the loss of catalytic potential. Unlike P1, the catalytic site P2 in PDX1.2 appears to be undisturbed except for the mutation of the key residue K166 to Q168.

Based on the structural analysis of overall shape and the comparison of areas around the catalytic sites between PDX1.2 and PDX1.3, there is no obvious benefit in keeping such a defective homolog and there is no clear mechanism about how PDX1.2 can positively regulate PDX1.3. We did not find any significant differences in the interfaces between subunits (Fig. S18). Hydrophobicity plots of PDX1.2 and PDX1.3 did not show any difference between the positions of polar and hydrophobic residues (Fig. S19). Molecular modeling using *Rosetta* showed no obvious impact of PDX1.2 on the stability of the complex. Surprisingly, the *Rosetta* calculation estimated that PDX1.3 packs more tightly than PDX1.2 and has more favorable sum of interactions and interface stablization (Table S2). Therefore, other regulatory contributions that cannot be captured by current computational methods are likely present (e.g. long-range surface electrostatics changes).

Interestingly, the most stunning differences were found when comparing the electrostatic surface potentials between the two (Fig. 4b). The surface potential of outer and inner regions of the PDX1.2 12mer, which spans the P1 and P2 sites, appears inverted relative to PDX1.3, where negative charges replace the positively charged regions. The difference in charge based on the structures was also consistent with the apparent lower stability of PDX1.3 than PDX1.2 in native MS as mentioned earlier. The positive electrostatic surface potential of PDX1.3 could induce charging and unfolding in electrospray, giving rise to significant dissociation in the native MS spectra.

## Discussion

Pseudoenzymes represent a largely uncharted territory with much to learn about the additional layer of regulatory control they impose in nature. They are now known for allosteric regulation and scaffolding through conserved binding interfaces or domains like their active enzyme homologs^5^. Yet, technical challenges exist in probing such pesudoenzyme-enzyme interactions due to extremely high structural similarity, as exemplified by the PDX1.2-PDX1.3 pseudoenzyme-enzyme pair detailed in this work.

A single particle cryo-EM approach was used to solve the structure of PDX1.2, a protein known not to crystallize. We found that the PDX1.2 fold closely mimics the fold of PDX1.3, a common trend observed for pseudoenzymes involved in allosteric regulation of their catalytic counterparts. PDX1.2 also displays an altered catalytic site P1 which appears perturbed in concordance with its lack of activity, while the P2 site remains largely unchanged. In the studies described herein, we were able to accurately control the stoichiometry of hetero-complexes using cell-free expression and characterize them using native MS. To our surprise, the catalysis can be tuned by shifting pseudoenzyme-enzyme stoichiometry, a discovery that was not reported previously for any known pseudoenzyme-enzyme pair today.

Additionally, the tunable nature of PDX1.2 and PDX1.3 co-complexes strongly disfavors the previously proposed assembly model that promoted interaction through homomeric 6mer rings (Fig.1a). Our native MS and SID data demonstrated that PDX1.2 co-assembly with its catalytic partner PDX1.3 is largely based on stochastic subunit incorporation at different locations but with some degree of symmetry (Fig.5a). Driven by the molar ratios of individual components during co-expression conditions, the entire range of all possible stoichiometry combinations (from 12:0 to 0:12) has been recorded.

**Fig.5.**
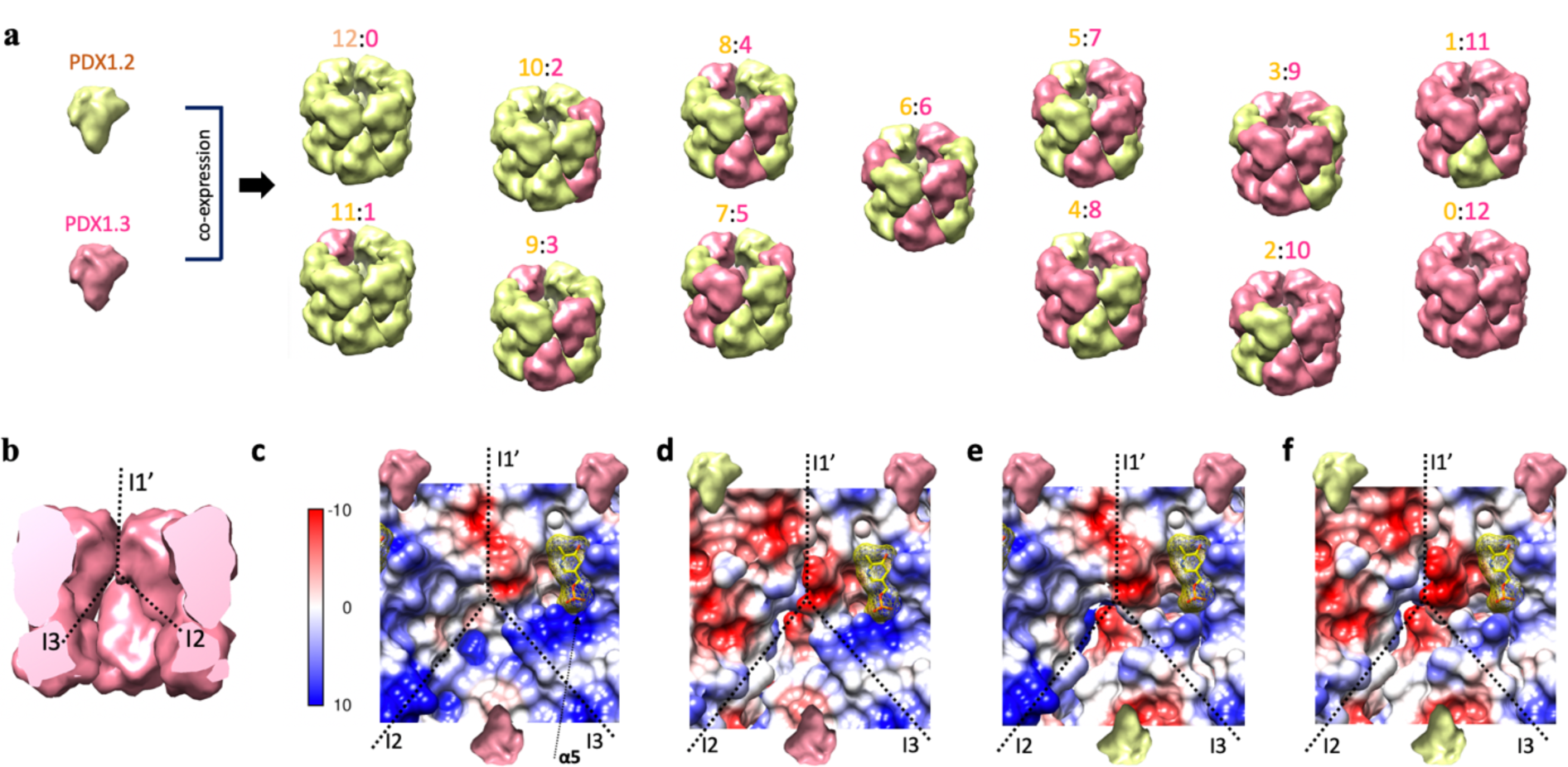
Heteromeric assembly mechanism and the proposed electrostatic reorganization around the P2 catalytic site. **a.** Revised model of PDX1.2 and PDX1.3 co-assembly. Each subclass is annotated as 12:0, 11:1, 10:2, 9:3 and et cetera, where the integers correspond to the number of PDX1.2 subunits relative to number of PDX1.3 subunits. Only one representative combination per class is shown while the total number of possible combinations is variable based on stoichiometry, symmetry operations and lateral favorability in the rings. **b.** The potential impact of PDX1.2 on PDX1.3 activity can be through I1’and I3 interfaces, surrounding the catalytic site P2 in each monomer. **c.** The electrostatic surface potential map of the area surrounding the P2 site in PDX1.3 homo-complex. The yellow molecule represents PLP. The models (**d-f**) were constructed by replacing PDX1.3 monomers with the cryo-EM structure of PDX1.2 monomers. Note the significantly changed electrostatic potential on the upper left of the P2 site in (**d**).

Another intriguing finding is the inverted surface charge of PDX1.2 compared to PDX1.3. It has been shown that surface charge-charge interactions can be redesigned to increase protein stability^31^. As PDX1.2 expression is induced under heat stress, the switch in the electrostatic surface potential might serve as a heat adaption mechanism. Our proposed structural models (Fig.5b-f) could help explain why the defective homolog PDX1.2 is preserved among plants. The catalytic site P1 is buried inside the center of the individual PDX1.3 subunit, and it is therefore unlikely to be impacted by PDX1.2. Instead, in concordance with the observed mechanism of assembly and amino acid conservation, we hypothesize that the most profound effect would be on the catalytic site P2. Site P2 is located at the edge of the PDX1.3 subunit and shares two subunit-subunit interfaces: side-by-side **I1’** within the hexamer ring and bottom-bottom **I3** at the ring-ring interface (Fig.5b). We constructed several PDX1.2-PDX1.3 assembly models around the catalytic site P2 of PDX1.3 by swapping subunits in the aligned homo-12mer structures (Fig.5c- f). Lateral PDX1.2 incorporation can create a local supercharging effect on the **ß4-α4** loop, which could potentially act on the nitrogen atom of its pyridoxine ring (Fig.5d). This supercharging effect could promote imine bond breaking between PLP and K166, help in the translocation and assist with the exit of the final product and thus increase the rate of reaction turnover. The supercharging of the same loop could also be possible when PDX1.2 shares a bottom-bottom interface (Fig.5e). The positioning of PDX1.2 on the bottom also slightly weakens the charge state around the phosphate binding site in **α4**. Both side and bottom consequent positioning of PDX1.2 (Fig.5f) could explain enhanced PDX1.3 activity in the hetero-complex and the benefit of having a pseudoenzyme that hetero-associates with active PDX1.3 in the 12mer.

While our current data cannot determine precisely which interfaces in the hetero-interaction are favored, the constructed models strongly suggest the benefit of such interactions for enzyme activity. Given the inverted electrostatic surface potentials of the two proteins, attractive forces between opposite charges may favor the symmetry in such hetero-interactions. We thus propose that PDX1.2 and PDX1.3 monomers are placed in an alternating manner within the hetero-complex so that the hetero-interactions can be maximized. This is supported by our observation that homo-subcomplexes were not released in SID for most hetero-complexes (Fig. S15) – meaning the same protomers do not prefer to cluster within the 12mer. Such positioning strategy empowers a simple, tunable and gentle approach to accelerate vitamin B_6_ production by active enzymes such as PDX1.3 and PDX1.1.

Although the monomer positions within individual hetero-complexes cannot be structurally resolved by either cryo-EM or X-ray crystallography, native MS provided accurate determination of stoichiometry and SID revealed the lateral symmetry in the subunit arrangement. Ongoing development in native MS instrumentation and the gas-phase protein behaviors would allow more structural information to be extracted from such experiments in the near future. Our study shows that complementing high resolution structural biology techniques such as cryo-EM with native MS offers new ways to understand heterogenous pseudoenzyme-enzyme interactions that cannot be elucidated by single techniques alone.

## Methods

### Plasmid construction

Gene sequences of PDX1.2 and PDX1.3 were sourced as described previously^19^. The clones were amplified by PCR to include the purification tag and then sub-cloned using Gibson Assembly in the designated vector pEU for the cell-free protein synthesis^19,32^. All reagents for PCR and Gibson reactions such as the Q5 Hot Start High-Fidelity 2X Master Mix and Gibson Assembly Master Mix were acquired from New England Biolabs. DNA primers were from ThermoFisher Scientific. For expression of PDX1.2 and PDX1.3 proteins, two vectors were constructed: pEU_3XFLAG_PDX1.2 and pEU_3XFLAG_PDX1.3, where both genes were fused with the 3XFLAG purification tag on their N-terminus (the sequences of synthesized proteins are in Fig. S1). The sequences of all plasmids used in this study were verified by Sanger sequencing (MCLAB).

### Cell-free expression/co-expression and purification of PDX proteins

Protein synthesis was carried out using Wheat Germ Protein Research Kits WEPRO7240 from CellFree Sciences. For co-expression, stock plasmids were prepared containing various amounts of pEU_3XFLAG_PDX1.2 and pEU_3XFLAG_PDX1.3. For example, in the case of co-expression at 9:1: 9 μl of 1 μg/μl pEU_3XFLAG_PDX1.2 was mixed with 1 μl of 1 μg/μl pEU_3XFLAG_PDX1.3 before the transcription step. The MINI-scale protein synthesis in the presence of fluorophore-charged lysine tRNA was conducted with the following conditions: (i) for transcription, 1 μl of 1 μg/μl stock plasmid was mixed with 9 μl of Transcription premix solution (1x Transcription buffer, 2.5 mM NTP, 8 U/μl SP6 polymerase, 8 U/μl RNase inhibitor) and left to incubate at 37°C for 4 hours, (ii) for translation, 2 μl of the resulting mRNA was mixed with 1 μl of FluoroTect Green_Lys_ (Promega) and 3 μl of the wheat germ extract and then transferred under 50 μl of the translation buffer (1x SUB-AMIX SGC) within a 96-well half-area plate. The translations were conducted overnight at 15°C and away from light in the vibration-free setting on an Eppendorf ThermoMixer C.

The MAXI-scale protein synthesis was performed using the robotic system - Protemist DT II from CellFree Sciences. All steps were performed at 4°C and at 140 rpm. The buffer exchange step of the crude extract (in order to remove dithiothreitol, DTT) was skipped. A total of 600 μl of ANTI-M2 affinity resin (50% suspension, pre-equilibrated in 1xTBS buffer, Sigma) was used per each 6 ml translation reaction. Protein binding to the resin was carried out for 1 hour. The resin was subsequently washed three times with 2.5 ml of 1xTBS buffer for 5 min each time. The protein products were eluted twice with the 1 ml Elution Buffer (1xTBS buffer, 100 μg/ml 3XFLAG peptide, 4 mM DTT) each time for 30 min. Proteins were further concentrated and buffer-exchanged to the Storage Buffer (1xTBS with 4 mM DTT) using 30 kDa Amicon Ultra-2 centrifugal filter units. The concentrated samples were flash-frozen in liquid nitrogen in 10 μl aliquots and then transferred to -80°C for storage.

General PAGE electrophoresis conditions employed in this work followed standard procedures. Bioanalyzer runs were performed with the Agilent Bioanalyzer 2100 system using the P80 kit according to the manufacturer’s guidelines.

### Peptide mapping

4 µg of protein was diluted in 50% trifluoroethanol and further incubated at 60 °C for 2 h at 800 rpm. Then 2 mM DTT was added and incubated at 37 °C for 1 h with shaking. The samples were diluted in 200 µL 25 mM ammonium bicarbonate, digested with 0.1 µg trypsin at 37 °C for 3 h with shaking. Both samples were dried down and re-suspended at 0.05 µg/µL in water for LCMS analysis. 5 µL peptide solution was loaded on a Waters M-Class NanoAcquity LC equipped with C18 reversed phase column (packed in-house, length 70 cm, 75 µm ID, 3µm Jupiter particle from Phenomenex). Peptides were separated using water/acetonitrile mobile phases with 0.1% formic acid over a gradient of 2h (ramping acetonitrile gradient 1-40%) at a flow rate of 300 nL/min. Mass spectra of eluting peptides were collected on a Thermo Q-Exactive Orbitrap. Resolution of 35k and 17.5k (at *m/z* 200) were used for MS^1^ and MS^2^, respectively. The top 12 precursors were selected for higher energy collision (HCD) at 30% normalized collision energy. A dynamic exclusion of 30s was used. Singly charged or charge unassigned precursors were excluded. LCMS data were analyzed with Byonic with a semi-tryptic search of the *Arabidopsis thaliana* protein database (source: Arabidopsis Information Resource), including common contaminants. The 3XFLAG purification tag sequence was manually appended to the original PDX1.2 and 1.3 sequences (AT3G16050.1 and AT5G01410.1). Precursor and fragment mass tolerances were set to 7 and 10 ppm. Protein N-terminal acetylation, pyroglutamic acid (Q), and oxidation (H, M, W) were included as dynamic modifications. Raw data and processing results are deposited in MassIVE (massive.ucsd.edu) with accession MSV000085233.

### Intact mass determination by LCMS

0.2-0.3 µg of protein diluted in water at 0.1 µg/µL were loaded on a Waters NanoAcquity LC equipped with a C2 reversed phase column (packed in-house, length 70 cm, 75 µm ID, 3µm C2 particle). Intact proteins were separated using water/acetonitrile mobile phases with 0.1% formic acid over a gradient of 1 h (ramping acetonitrile gradient 10-50%) at a flow rate of 300 nL/min. Mass spectra of eluting proteins were collected on a Thermo VelosOrbitrap Elite. MS^1^ spectra were acquired at 120k resolution (at *m/z* 200) and were averaged over 8 microscans. ProMex^33^ was used to de-convolute the data and obtain the intact mass of the proteins. LCMS data were aggregated across retention time and binned into unit mass (sum intensities within 1 Da) to generate the intact mass plot in Fig. S5. Raw data and processing results are deposited in MassIVE (massive.ucsd.edu) with accession MSV000085233.

### Native Mass Spectrometry (native MS)

Purified PDX complexes at 0.2-0.4 mg/mL concentrations (10 µL each in 1xTBS with 4 mM DTT) were buffer exchanged into 200 mM ammonium acetate (pH 6.7) using Zeba 75 µL microspin columns from Thermofisher. Proteins were then loaded into pulled glass capillaries (GlassTip™, part number BG12692N20, Scientific Instrument Services, Inc., Ringoes, NJ) with gel loading pipet tips. A platinum wire connected to the electrospray voltage was inserted into the capillary and was in contact with the protein solution. 1 kV was applied at the capillary to sustain the nanoelectrospray on a Thermo Q-Exactive UHMR mass spectrometer. The source was kept at 200 °C, with the S-lens set to 100%. In-source fragmentation of 100 V and in-source trapping of 50 V were used for optimum de-solvation to measure the mass of the complexes without extensive gas-phase dissociation. Even with in-source fragmentation/trapping voltages off, a significant amount of PDX1.3 monomers were still observed, implying that the PDX1.3 monomers were not exclusively generated by the desolvation conditions. HCD gas flow was set to 2. For each final spectrum, 500 microscans were averaged. Mass calibration was performed using cesium iodide clusters. To obtain the mass distribution of the PDX 12mer complexes, mass spectra were processed using UniDec v2.7^31^. Original data within m/z 8000-11000 were extracted with curved baseline subtraction, Gaussian smoothing 2.0, and binned every 3 m/z. Deconvolution was restricted to sampling mass every 10 Da with peak FWHM of 8.

### Activity assay

For quantitative PLP detection, we used Enzymatic Vitamin B6 assay from A/C diagnostics (http://www.vitaminb6assay.com/). From an initial set of experiments using active enzyme PDX1.3, we determined that 5-10 ug/ml was the optimal protein concentration for conducting this assay (in place of the original design of using blood plasma) as this range results in the OD675 signal falling within the linear PLP detection range of 0-200 nmol/L (data not shown).

To compare relative enzymatic activities, all PDX samples (individual or co-expressed) were first diluted to 12.5 ug/ml concentration in TBS buffer. Then 5 ul of 12.5 ug/ml protein were combined with 5 ul of 2XAssay buffer (50 mM Tris, pH 8.0, 1 mM ribose 5-phosphate, 2 mM D-glyceraldehyde 3-phosphate, 20 mM ammonium sulfate) and incubated away from light for 30 min at 30°C in the Eppendor thermocycler. For the calibration curve, PLP (Sigma, Cat # 82870) was diluted in a serial manner to 400 nM, 200 nM, 100 nM, 50 nM, 25 nM concentration in TBS buffer, further combined with an equal volume of 2XAssay buffer and incubated away from light for 30 min at 30°C along with PDX samples for consistency.

To quantify the amount of PLP in the mixture, 2.5 ul of each sample were further added to the 37.5 ul of Working Binding Assay buffer (with apo-enzyme), provided by the assay kit from A/C diagnostics and placed in the Corning NBS 384-well microplate. The mixture was incubated at 37°C for 30 min at 750 rpm on the Eppendorf Thermomixer C away from light. Then 20 ul of Working Assay buffer (from the kit) was added to the mixture, and incubation continued for additional 20 min. This was followed by the addition of 6.25 ul of Chromogen RI and 3.75 ul of Chromogen RII and 10 min incubation. The color development was measured at 675 nm using the Infinite M200 PRO microplate reader by Tecan. For accuracy, all samples and calibration standards were assayed in triplicate.

### Surface induced dissocation (SID)

Purified PDX complexes were buffer exchanged into 200 mM ammonium acetate (pH 6.7) using the same protocol described for native MS. Then the protein solution was mixed 1:1 with 100 mM triethylammonium acetate (TEAA), and loaded into in-house pulled glass capillaries (Sutter Instruments P-97 micropipet puller,with glass capillary catalog BF100-78-10). The same instrument conditions were used as native MS but on a modified Thermo Q-Exactive UHMR mass spectrometer as described in a previous report^23^. Briefly, a custom made SID cell was inserted after the quadrupole and before the C-trap, replacing the original hexapole in the commercial configuration. PDX complexes were mass-isolated with a 30-35 *m/z* window for the 35+ charge state, and steered towards the surface under a controlled accelearation voltage of 160 V (5.6 keV). Product ions after the surface impact were collected in the C-trap and mass analyzed in the Orbitrap. The electrospray voltage was set to 1.2-1.3 kV, S-lens set at 100%, in-source trapping set at -100 V, and HCD gas flow set to 4. Spectra were collected at a resolution of 6,250 (at *m/z* of 400), with an injection time of 1 s and averaged over 200-500 scans. Mass calibration was performed using cesium iodide clusters. SID tuning voltages are listed in Table S3. SID spectra were deconvoluted using UniDec v4.2.1^34^. Mass was sampled every 10 Da with the peak FWHM set to 6, charge state smooth set at 2 and point smooth set to 1. Experimental peaks were fit to all possible combinations of PDX1.2/PDX1.3 oligomers using the mass list option in UniDec. A DScore of 30% and a peak threshold of 0.01 were used to filter the deconvolution results.

In the deconvoluted results (Fig. S15), peak areas were used for calculation of relative abundances of different 6mer species. The peak areas of the 6mers from both the major pathway (horizontal cleavage in Fig. 3c) and the minor pathways (vertical cleavage in Fig. 3c or secondary dissociation) cannot be directly distinguished by mass. We assume that the minor pathways generate different types of products with similar abundances. Thus, the peak areas of the 6mers from the minor pathways can be estimated by the median peak area of the detected 4mers, 5mers, 7mers, and 8mers in the same SID spectrum. For precursor 9:3, 10:2, and 11:1, only 6mers were consistently detected and the medians were set to 0. The peak areas of the 6mers from the major pathway were calculated by subtracting the contributions of minor pathways from the experimtal peak areas. If the values are negative, they are reset to 0. The “trimmed” abundances were normalized and plotted in Fig. 3d and Fig.S15c. The relative abundances calculated without “trimming” are shown in Fig. S15d.

### Cryo-EM data acquisition and processing

Quantifoil grids (658-300-AU, Ted Pella) were first glow-discharged at 15 mA for 1 min using PELCO easiGlow (Ted Pella). The grids were then transferred to a Leica EM GP the plunge-freezer, brought to 85% humidity at 25°C and 3 μl of protein sample (0.2 mg/ml, in TBS buffer with 4 mM DTT) was pipetted on the carbon side. The grid with sample was then blotted for 3 s, plunge-frozen in the liquid ethane and transferred to liquid nitrogen. The frozen samples were imaged on a 300 keV FEI Titan Krios at Pacific Northwest Center for Cryo-EM (https://pncc.labworks.org) equipped with a Gatan K2 summit direct electron detector. The data was collected using automated acquisition software SerialEM^35^. For PDX1.2 sample, 8,106 -resolution movies with 60 frames each at a dosage of 1.5 electrons per Å^2^ per frame and pixel size of 0.253 Å were collected. For PDXcoexp, 6,904 super-resolution movies with 50 frames each at a dosage of 2 electrons per Å ^2^ per frame and pixel size of 0.253 Å were acquired. All data were processed using cryoSPARC v2 software^36^ and were corrected for full frame motion using MotionCor2^37^ and CTF estimated using CTFFIND4^38^. For PDX1.2 sample, a total of 787,982 particles were identified from template-based auto-picking algorithm. Three rounds of reference-free 2D classification narrowed the pool to 265,224 particles. These were used to generate the ab initio 3D map as a reference. Homogenous refinement in cryoSPARC v2 with D6 symmetry produced a final 3.2 Å reconstruction. For PDXcoexp sample, a total of 510,660 particles were identified from template-based auto-picking algorithm. Three rounds of reference-free 2D classification narrowed the pool to 286,642 particles. These were used to generate the ab initio 3D map as a reference. Homogenous refinement in cryosparc v2 with D6 symmetry produced 3.2 Å reconstruction while non-uniform refinement (BETA) yielded the 3.16 Å map. Local resolution calculations on the final maps were performed in Relion 3^39^.

### Atomic modeling

An initial homology model of PDX1.2 monomer was generated using HHPRED^40^ and MODELLER^41^ using comparative modeling against known structures for PDX from other organisms (2NV1, 5LNR, 4JDY, 2YZR and 3O07). The Cryo-EM map and the aligned initial model for a single subunit were imported to PHENIX^42^, docked using *Phenix.DockInMap*, and the density region surrounding the monomer unit mapped out via *Phenix.MapBox*. Several cycles of *Phenix.RealspaceRefine* were carried out to refine the atomic model for the individual monomer. Map symmetry parameters were then applied on the real space refined monomer model using *Phenix.ApplyNCSoperators* to generate the 12-mer, which underwent additional rounds of *Phenix.RealspaceRefine*. Final refinement statistics and validation scores are presented in Table S1. The final models and maps were uploaded to the PDB and EMDB databases under the accession numbers: **6PCJ** and **EMD-20302** for PDX1.2; and **6PCN** and **EMD- 20303** for PDXcoexp. Multiscale models, structural comparisons, structure-based alignments and computing of electrostatic potentials were conducted in Chimera^43^. Protein-protein interactions analysis was perfomed by PDBsum server^44^.

### Rosetta calculations

*Rosetta3*^45^ was used to characterize the features of PDX proteins (Table S2). *FastRelax*^46^ with the latest score function (*ref2015_cart*) was perfomed before analyzing full atom energy values. To better pack protein structure, we used *dualspace*^47^ when we relaxed structures. Among 100 decoys (“nstruct”) per each relaxation, we used the top 10% decoys in terms of lower (“better”) total score values. For interface analysis and ligand binding energy calculation, we used the lowest scored decoy. For interface analysis and packing status, we used *InterfaceAnalyzer*^48^ and *RosettaHoles*^49^ respectively. To report feature values and analyze effectively, we used *RosettaScripts*^50^.

## Acknowledgements

This work was supported in part by DOE-BER Mesoscale to Molecules Bioimaging Project FWP# 66382 and EMSL Strategic Science Area Projects 50188, 50165, and 50427. A portion of this research was supported by NIH grant U24GM129547 and performed at the PNCC at OHSU and accessed through EMSL (grid.436923.9), a DOE Office of Science User Facility sponsored by the Office of Biological and Environmental Research. CD, ZLV, and VHW acknowledge the support by the National Institutes of Health Grant P41 GM128577. We thank Carrie Nicora, Karl Weitz, Anil Shukla, Ronald Moore, and Rui Zhao for assistance with the LCMS experiments and Harry Scott for expert data collection at PNCC. We thank also Matthew Monroe for helping with MassIVE data deposition.

## Author contributions

JEE, IVN and MZ devised experiments for the study. IVN performed molecular biology, cell-free expression, cryoEM data processing and atomic modeling. MZ designed the all MS, native MS, and SID experiments, and performed most of the related data collection and analysis. CD performed the SID data collection and analysis. ZLV assisted with the SID experiments. JS assisted with the native MS setup at EMSL. DNK had performed *Rosetta* calcualtions. MP and HH provided clones of PDX and assisted with analysis of the resulting proteins. MP also performed a portion of cell-free experiments. IVN, JEE and MZ wrote initial manuscript draft but all authors contributed to writing the manuscript and approval of the final version.

## Competing Interests

The authors declare no competing interests or conflicts of interest.

## Notes

### Competing Interest Statement

The authors have declared no competing interest.

### Summary of Updates

New data has been incorporated

## References

1. Murphy, J. M., Mace, P. D. & Eyers, P. A. Live and let die: insights into pseudoenzyme mechanisms from structure. Curr Opin Struct Biol 47, 95–104, doi:10.1016/j.sbi.2017.07.004 (2017).

2. Yu, J. W., Jeffrey, P. D. & Shi, Y. Mechanism of procaspase-8 activation by c-FLIPL. Proc Natl Acad Sci U S A 106, 8169–8174, doi:10.1073/pnas.0812453106 (2009).

3. Murphy, J. M. et al. The pseudokinase MLKL mediates necroptosis via a molecular switch mechanism. Immunity 39, 443–453, doi:10.1016/j.immuni.2013.06.018 (2013).

4. Schimpl, M. et al. Human YKL-39 is a pseudo-chitinase with retained chitooligosaccharide-binding properties. Biochem J 446, 149–157, doi:10.1042/BJ20120377 (2012).

5. Ribeiro, A.J. M. et al. Emerging concepts in pseudoenzyme classification, evolution, and signaling. Sci Signal 12, doi:10.1126/scisignal.aat9797 (2019).

6. Parra, M., Stahl, S. & Hellmann, H. Vitamin B(6) and Its Role in Cell Metabolism and Physiology. Cells 7, doi:10.3390/cells7070084 (2018).

7. Mooney, S., Leuendorf, J. E., Hendrickson, C. & Hellmann, H. Vitamin B6: a long known compound of surprising complexity. Molecules 14, 329–351, doi:10.3390/molecules14010329 (2009).

8. Hsu, C. C. et al. Role of vitamin B6 status on antioxidant defenses, glutathione, and related enzyme activities in mice with homocysteine-induced oxidative stress. Food Nutr Res 59, 25702, doi:10.3402/fnr.v59.25702 (2015).

9. Zhang, Y. F. et al. The de novo Biosynthesis of Vitamin B6 Is Required for Disease Resistance Against Botrytis cinerea in Tomato. Mol Plant Microbe In 27, 688–699, doi:10.1094/Mpmi-01-14-0020-R (2014).

10. Chandrasekaran, M., Paramasivan, M. & Chun, S. C. Bacillus subtilis CBR05 induces Vitamin B6 biosynthesis in tomato through the de novo pathway in contributing disease resistance against Xanthomonas campestris pv. vesicatoria. Sci Rep-Uk 9, doi:10.1038/s41598-019-41888-6 (2019).

11. Tambasco-Studart, M. et al. Vitamin B6 biosynthesis in higher plants. Proc Natl Acad Sci U S A 102, 13687–13692, doi:10.1073/pnas.0506228102 (2005).

12. Leuendorf, J. E., Mooney, S. L., Chen, L. & Hellmann, H. A. Arabidopsis thaliana PDX1.2 is critical for embryo development and heat shock tolerance. Planta 240, 137–146, doi:10.1007/s00425-014-2069-3 (2014).

13. Moccand, C. et al. The pseudoenzyme PDX1.2 boosts vitamin B6 biosynthesis under heat and oxidative stress in Arabidopsis. J Biol Chem 289, 8203–8216, doi:10.1074/jbc.M113.540526 (2014).

14. Denslow, S. A., Rueschhoff, E. E. & Daub, M. E. Regulation of the Arabidopsis thaliana vitamin B6 biosynthesis genes by abiotic stress. Plant Physiol Biochem 45, 152–161, doi:10.1016/j.plaphy.2007.01.007 (2007).

15. Leuendorf, J. E., Osorio, S., Szewczyk, A., Fernie, A. R. & Hellmann, H. Complex assembly and metabolic profiling of Arabidopsis thaliana plants overexpressing vitamin B(6) biosynthesis proteins. Mol Plant 3, 890–903, doi:10.1093/mp/ssq041 (2010).

16. Robinson, G. C. et al. Crystal structure of the pseudoenzyme PDX1.2 in complex with its cognate enzyme PDX1.3: a total eclipse. Acta Crystallogr D Struct Biol 75, 400–415, doi:10.1107/S2059798319002912 (2019).

17. Madin, K., Sawasaki, T., Ogasawara, T. & Endo, Y. A highly efficient and robust cell-free protein synthesis system prepared from wheat embryos: plants apparently contain a suicide system directed at ribosomes. Proc Natl Acad Sci U S A 97, 559–564, doi:10.1073/pnas.97.2.559 (2000).

18. Sawasaki, T. et al. A bilayer cell-free protein synthesis system for high-throughput screening of gene products. FEBS Lett 514, 102–105 (2002).

19. Novikova, I. V. et al. Protein structural biology using cell-free platform from wheat germ. Adv Struct Chem Imaging 4, 13, doi:10.1186/s40679-018-0062-9 (2018).

20. Titiz, O. et al. PDX1 is essential for vitamin B6 biosynthesis, development and stress tolerance in Arabidopsis. Plant J 48, 933–946, doi:10.1111/j.1365-313X.2006.02928.x (2006).

21. Wagner, S. et al. Analysis of the Arabidopsis rsr4-1/pdx1-3 mutant reveals the critical function of the PDX1 protein family in metabolism, development, and vitamin B6 biosynthesis. Plant Cell 18, 1722–1735, doi:10.1105/tpc.105.036269 (2006).

22. Mortensen, D. N. & Williams, E. R. Electrothermal supercharging of proteins in native MS: effects of protein isoelectric point, buffer, and nanoESI-emitter tip size. Analyst 141, 5598–5606, doi:10.1039/c6an01380e (2016).

23. Leney, A. C. Subunit pI can influence protein complex dissociation characteristics. J Am Soc Mass Spectr 30, 1389–1395, doi:10.1007/s13361-019-02198-3 (2019).

24. Hall, Z., Hernandez, H., Marsh, J. A., Teichmann, S. A. & Robinson, C. V. The role of salt bridges, charge density, and subunit flexibility in determining disassembly routes of protein complexes. Structure 21, 1325–1337, doi:10.1016/j.str.2013.06.004 (2013).

25. VanAernum, Z. L. et al. Surface-induced dissociation of noncovalent protein complexes in an extended mass range Orbitrap Mass Spectrometer. Anal Chem 91, 3611–3618, doi:10.1021/acs.analchem.8b05605 (2019).

26. Zhou, M., Dagan, S. & Wysocki, V. H. Impact of charge state on gas-phase behaviors of noncovalent protein complexes in collision induced dissociation and surface induced dissociation. Analyst 138, 1353–1362, doi:10.1039/c2an36525a (2013).

27. Stiving, A. Q. et al. Surface-induced dissociation: an effective method for characterization of protein quaternary structure. Anal Chem 91, 190–209, doi:10.1021/acs.analchem.8b05071 (2019).

28. Harvey, S. R. et al. Relative interfacial cleavage energetics of protein complexes revealed by surface collisions. Proc Natl Acad Sci U S A 116, 8143–8148, doi:10.1073/pnas.1817632116 (2019).

29. Rodrigues, M. J. et al. Lysine relay mechanism coordinates intermediate transfer in vitamin B6 biosynthesis. Nat Chem Biol 13, 290–294, doi:10.1038/nchembio.2273 (2017).

30. Robinson, G. C., Kaufmann, M., Roux, C. & Fitzpatrick, T. B. Structural definition of the lysine swing in Arabidopsis thaliana PDX1: Intermediate channeling facilitating vitamin B6 biosynthesis. Proc Natl Acad Sci U S A 113, E5821–E5829, doi:10.1073/pnas.1608125113 (2016).

31. Strickler, S. S. et al. Protein stability and surface electrostatics: a charged relationship. Biochemistry-Us 45, 2761–2766, doi:10.1021/bi0600143 (2006).

32. Sawasaki, T., Hasegawa, Y., Tsuchimochi, M., Kasahara, Y. & Endo, Y. Construction of an efficient expression vector for coupled transcription/translation in a wheat germ cell-free system. Nucleic Acids Symp Ser, 9–10 (2000).

33. Park, J. et al. Informed-Proteomics: open-source software package for top-down proteomics. Nat Methods 14, 909–914, doi:10.1038/nmeth.4388 (2017).

34. Marty, M. T. et al. Bayesian deconvolution of mass and ion mobility spectra: from binary interactions to polydisperse ensembles. Anal Chem 87, 4370–4376, doi:10.1021/acs.analchem.5b00140 (2015).

35. Mastronarde, D. N. Automated electron microscope tomography using robust prediction of specimen movements. J Struct Biol 152, 36–51, doi:10.1016/j.jsb.2005.07.007 (2005).

36. Punjani, A., Rubinstein, J. L., Fleet, D. J. & Brubaker, M. A. cryoSPARC: algorithms for rapid unsupervised cryo-EM structure determination. Nat Methods 14, 290–296, doi:10.1038/nmeth.4169 (2017).

37. Zheng, S. Q. et al. MotionCor2: anisotropic correction of beam-induced motion for improved cryoelectron microscopy. Nat Methods 14, 331–332, doi:10.1038/nmeth.4193 (2017).

38. Rohou, A. & Grigorieff, N. CTFFIND4: Fast and accurate defocus estimation from electron micrographs. J Struct Biol 192, 216–221, doi:10.1016/j.jsb.2015.08.008 (2015).

39. Zivanov, J. et al. New tools for automated high-resolution cryo-EM structure determination in RELION-3. Elife 7, doi:10.7554/eLife.42166 (2018).

40. Zimmermann, L. et al. A Completely Reimplemented MPI Bioinformatics Toolkit with a New HHpred Server at its Core. J Mol Biol 430, 2237–2243, doi:10.1016/j.jmb.2017.12.007 (2018).

41. Webb, B. & Sali, A. Protein Structure Modeling with MODELLER. Methods Mol Biol 1654, 39–54, doi:10.1007/978-1-4939-7231-9_4 (2017).

42. Adams, P. D. et al. The Phenix software for automated determination of macromolecular structures. Methods 55, 94–106, doi:10.1016/j.ymeth.2011.07.005 (2011).

43. Pettersen, E. F. et al. UCSF Chimera--a visualization system for exploratory research and analysis. J Comput Chem 25, 1605–1612, doi:10.1002/jcc.20084 (2004).

44. Laskowski, R. A., Jablonska, J., Pravda, L., Varekova, R. S. & Thornton, J. M. PDBsum: Structural summaries of PDB entries. Protein Sci 27, 129–134, doi:10.1002/pro.3289 (2018).

45. Leaver-Fay, A. et al. Rosetta3: An Object-Oriented Software Suite for the Simulation and Design of Macromolecules. Method Enzymol, 545–574, doi:10.1016/S0076-6879(11)87019-9 (2011).

46. Tyka, M. D. et al. Alternate states of proteins revealed by detailed energy landscape mapping. J Mol Biol 405, 607–618, doi:10.1016/j.jmb.2010.11.008 (2011).

47. Conway, P., Tyka, M. D., DiMaio, F., Konerding, D. E. & Baker, D. Relaxation of backbone bond geometry improves protein energy landscape modeling. Protein Sci 23, 47–55, doi:10.1002/pro.2389 (2014).

48. Stranges, P. B. & Kuhlman, B. A comparison of successful and failed protein interface designs highlights the challenges of designing buried hydrogen bonds. Protein Sci 22, 74–82, doi:10.1002/pro.2187 (2013).

49. Sheffler, W. & Baker, D. RosettaHoles: rapid assessment of protein core packing for structure prediction, refinement, design, and validation. Protein Sci 18, 229–239, doi:10.1002/pro.8 (2009).

50. Fleishman, S. J. et al. RosettaScripts: a scripting language interface to the Rosetta macromolecular modeling suite. PLoS One 6, e20161, doi:10.1371/journal.pone.0020161 (2011).

