## Supporting Info for "Tunable hetero-assembly of a plant pseudoenzyme-enzyme complex"

```
>PDX1.2
MDYKDHDGDYKDHDIDYKDDDDKLAGGGGSGGGGSADQAMTDQDQGAVTLYSGTAITDAKKNHPFSVKVG
LAQVLRGGAIVEVSSVNQAKLAESAGACSVIVSDPVRSRGGVRRMPDPVLIKEVKRAVSVPVMARARVGH
FVEAQILESLAVDYIDESEIISVADDDHFINKHNFRSPFICGCRDTGEALRRIREGAAMIRIQGDLTATG
NIAETVKNVRSLMGEVRVLNNMDDDEVFTFAKKISAPYDLVAQTKQMGRVPVVQFASGGITTPADAALMM
QLGCDGVFVGSEVFDGPDPFKKLRSIVQAVQHYNDPHVLAEMSSGLENAMESLNVRGDRIQDFGQGSV*

>PDX1.3
MDYKDHDGDYKDHDIDYKDDDDKLAGGGGSEGTGVVAVYGNGAITEAKKSPFSVKVGLAQMLRGGVIMDV
VNAEQARIAEEAGACAVMALERVPADIRAQGGVARMSDPQMIKEIKQAVTIPVMAKARIGHFVEAQILEA
IGIDYIDESEVLTLADEDHHINKHNFRIPFVCGCRNLGEALRRIREGAAMIRTKGEAGTGNIIEAVRHVR
SVNGDIRVLRNMDDDEVFTFAKKLAAPYDLVMQTKQLGRLPVVQFAAGGVATPADAALMMQLGCDGVFVG
SGIFKSGDPARRARAIVQAVTHYSDPEMLVEVSCGLGEAMVGINLNDEKVERFANRSE*
```

Fig. S1. Amino acid sequences of synthesized PDX1 proteins. The cleavable 3XFLAG sequence (used as a purification tag) is in red; the amino acid linker is in blue. Different linker lengths were employed for PDX1.2 and PDX1.3 to create a larger molecular weight difference between the two proteins to aid PAGE and native mass spectrometry analysis.

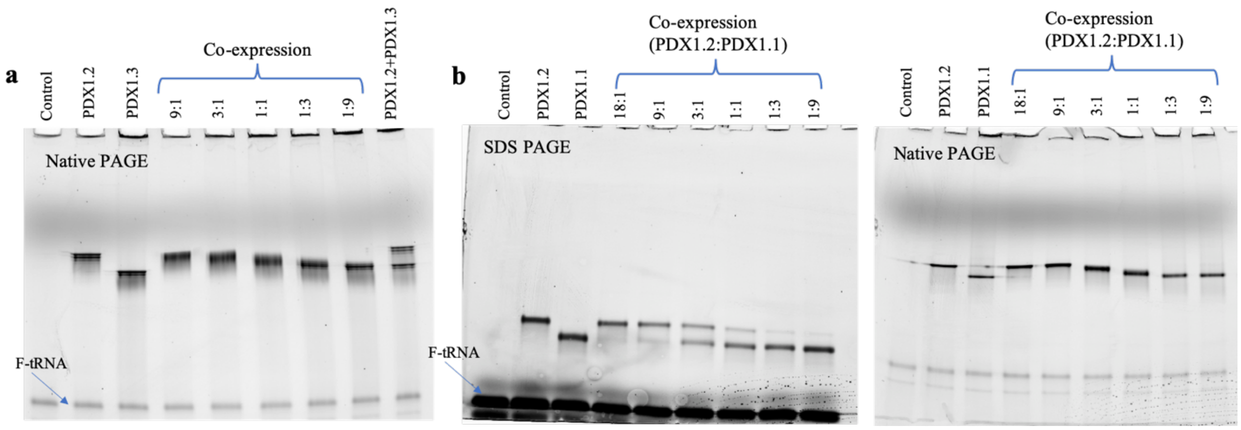


Fig.S2. Additional gel electrophoresis data for PDX1.2-PDX1.3 and PDX1.2-PDX1.1 pseudoenzyme-enzyme complexes. **a**. Native PAGE conducted on the same samples as in Fig.1c but after 48 hours incubation at 4°C. No observable degradation of complexes was detected compared to Figure 1. The last lane shows the individually expressed homomeric PDX1.2 and PDX1.3 proteins mixed post-translationally and incubated for 48 hours. The PDX1.2 and PDX1.3 remained homomeric in the mixture and no formation of intermixed complexes were observed as shown by distinct bands matching to the two individual proteins. **b**. PAGE data on the co-expression of PDX1.2 and PDX1.1 proteins in the crude wheat germ mixture. Co-expressed conditions are denoted as 18:1, 9:1, 3:1, 1:1, 1:3 and 1:9, where these numbers correspond to DNA template ratio of PDX1.2 to PDX1.1. PAGE data include a denaturing SDS-PAGE gel on the left and Native PAGE on the right. F-tRNA stands for fluorophore-labeled lysine-charged tRNA. Note that a similar trend in complexation is seen for PDX1.2-PDX1.3 pair, as shown in panel **(a)** and Fig.1c.

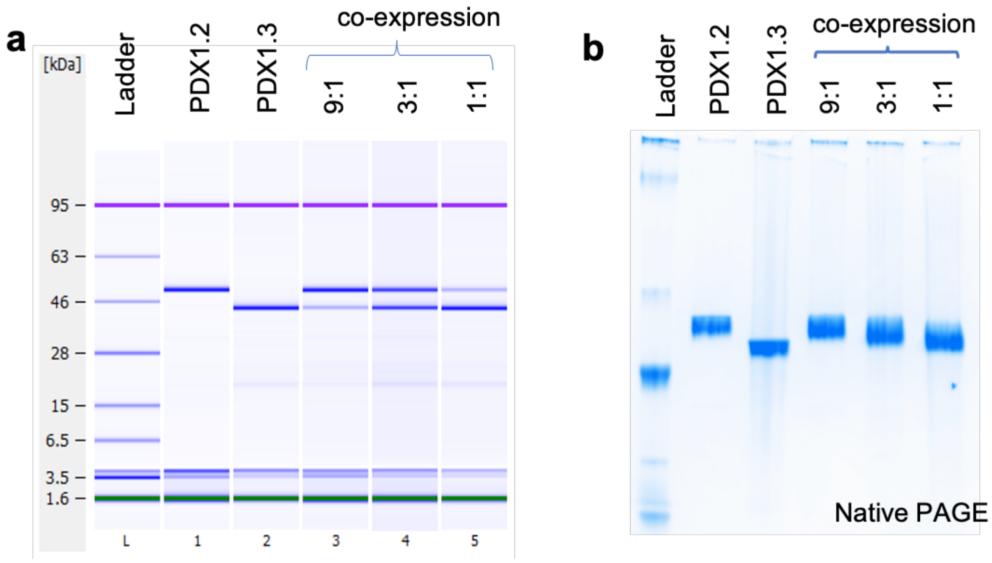


Fig. S3. Analysis of purified PDX complexes. **a**. Denaturing Bioanalyzer Protein 80 electrophoregram of purified PDX protein samples. **b**. Native PAGE. Ladder stands for NativeMark Protein Standard (ThermoFisher). Note that co-expressed samples have wider bands, consistent with Fig.1c.

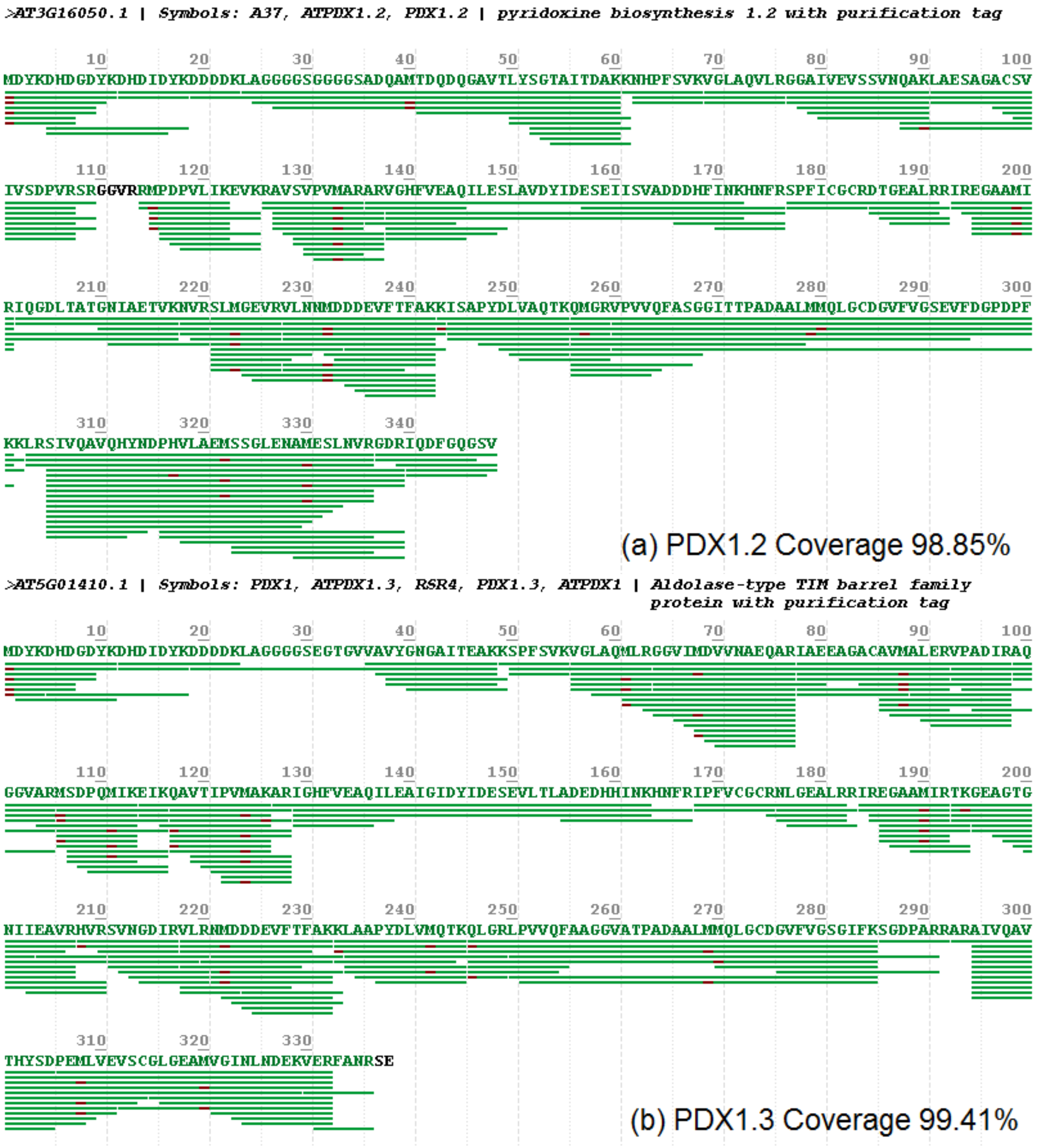


Fig. S4. Peptide mapping coverage map for cell-free expressed (a) PDX1.2 and (b) PDX1.3. Green horizontal lines along the protein sequences show the peptides detected. Protein residues colored in black are the regions where no peptides were detected (too short to be detected). Purple segments along the sequences are modified residues (primarily methionine oxidation and protein N-terminal oxidation), induced by the sample preparation.

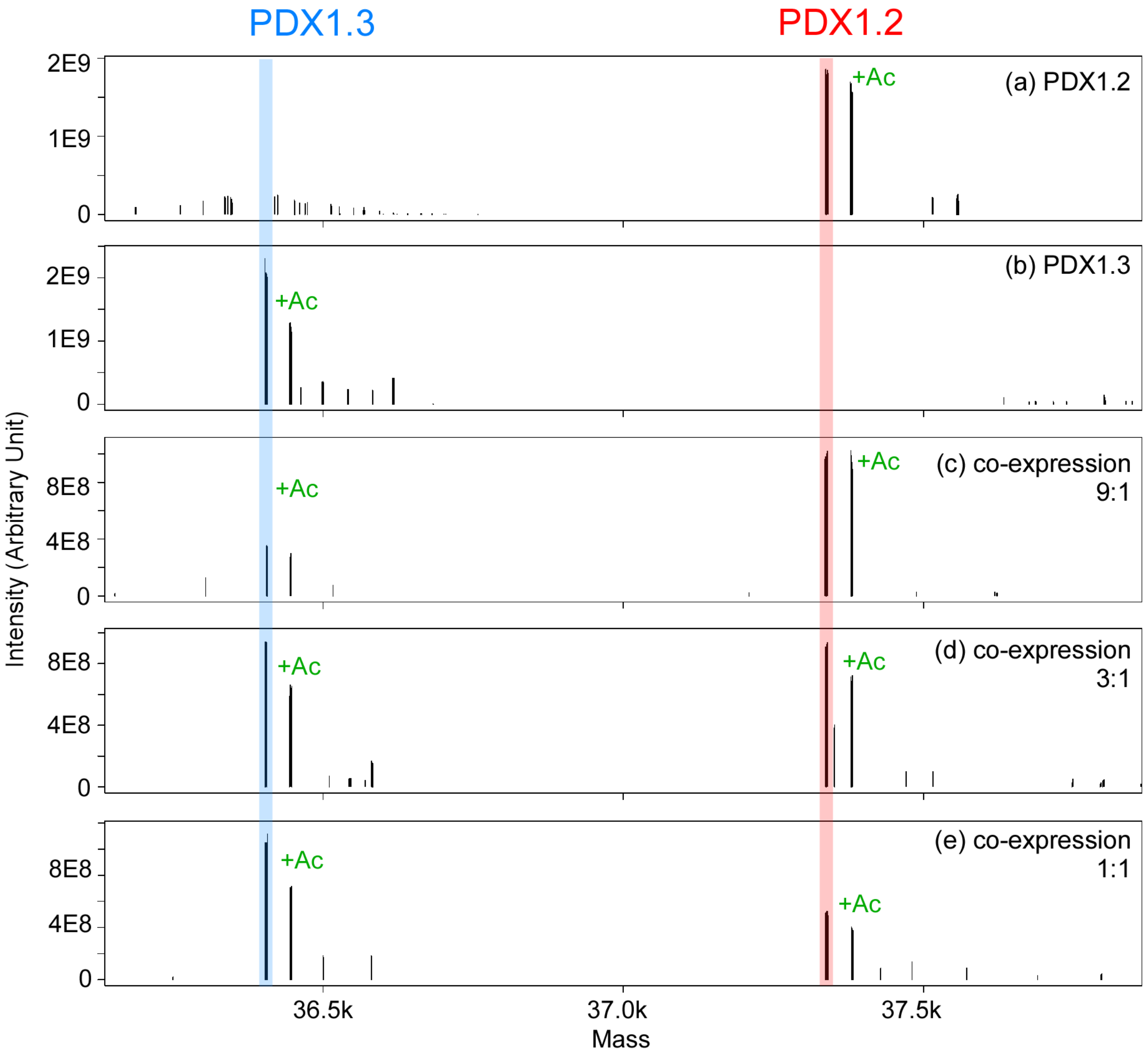


Fig. S5. Deconvoluted mass spectra of denatured intact PDX proteins for (a) PDX 1.2, (b) PDX 1.3, (c) co-expression 9:1, (d) co-expression 3:1, and (e) co-expression 1:1. About half of the protein show additional peak due to acetylation at the protein N-terminus. In all homomeric PDX1.2 and PDX1.3 samples and all three co-expressed samples, roughly half of the protein population was acetylated. These modifications, which are incorporated in the protein during its synthesis by ribosome-bound N-terminal acetyltransferases (NATs), are the most abundant, especially in higher eukaryotes *(Arnesen, T. Towards a functional understanding of protein. N-terminal acetylation. PLoS Biol 9, e1001074, doi:10.1371/journal.pbio.1001074 (2011))*. In general, such acetyl modifications are associated with various protein degradation and stabilization mechanisms inside the cell *(Hwang, C. S., Shemorry, A. & Varshavsky, A. N-terminal acetylation of cellular proteins creates specific degradation signals. Science 327, 973-977, doi:10.1126/science.1183147 (2010))*. Since no preferential acetylation profile was found across the samples, these modifications are likely irrelevant to PLP regulation itself.

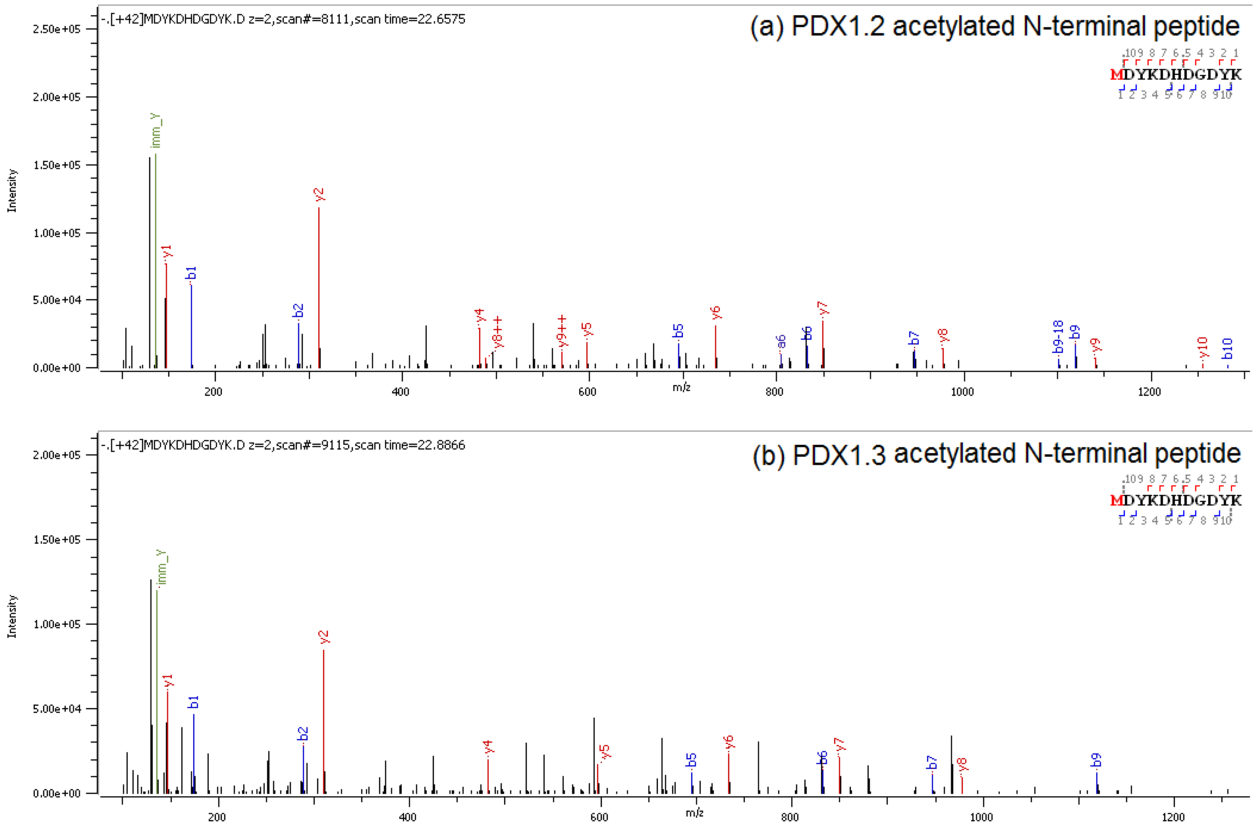


Fig. S6. Annotated $MS^2$ spectra of acetylated N-terminal peptide for (a) PDX1.2 and (b) PDX1.3. Fragmentation spectra suggest that acetylations are localized at the N termini of the proteins. In the previous study *(Moccand, C. et al. The pseudoenzyme PDX1.2 boosts vitamin B6 biosynthesis under heat and oxidative stress in Arabidopsis. J Biol Chem 289, 8203-8216, doi: 10.1074/jbc.M113.540526 (2014))*, *E. coli* expressed PDX1.2 was found to carry an acetyl group on the alanine (A5). Such modification was not detected in the wheat germ cell-free produced PDX1.2 and PDX1.3 shown here. N-terminal acetyl groups were detected and located on the first methionine residue, preceding the exogenous protein tag.

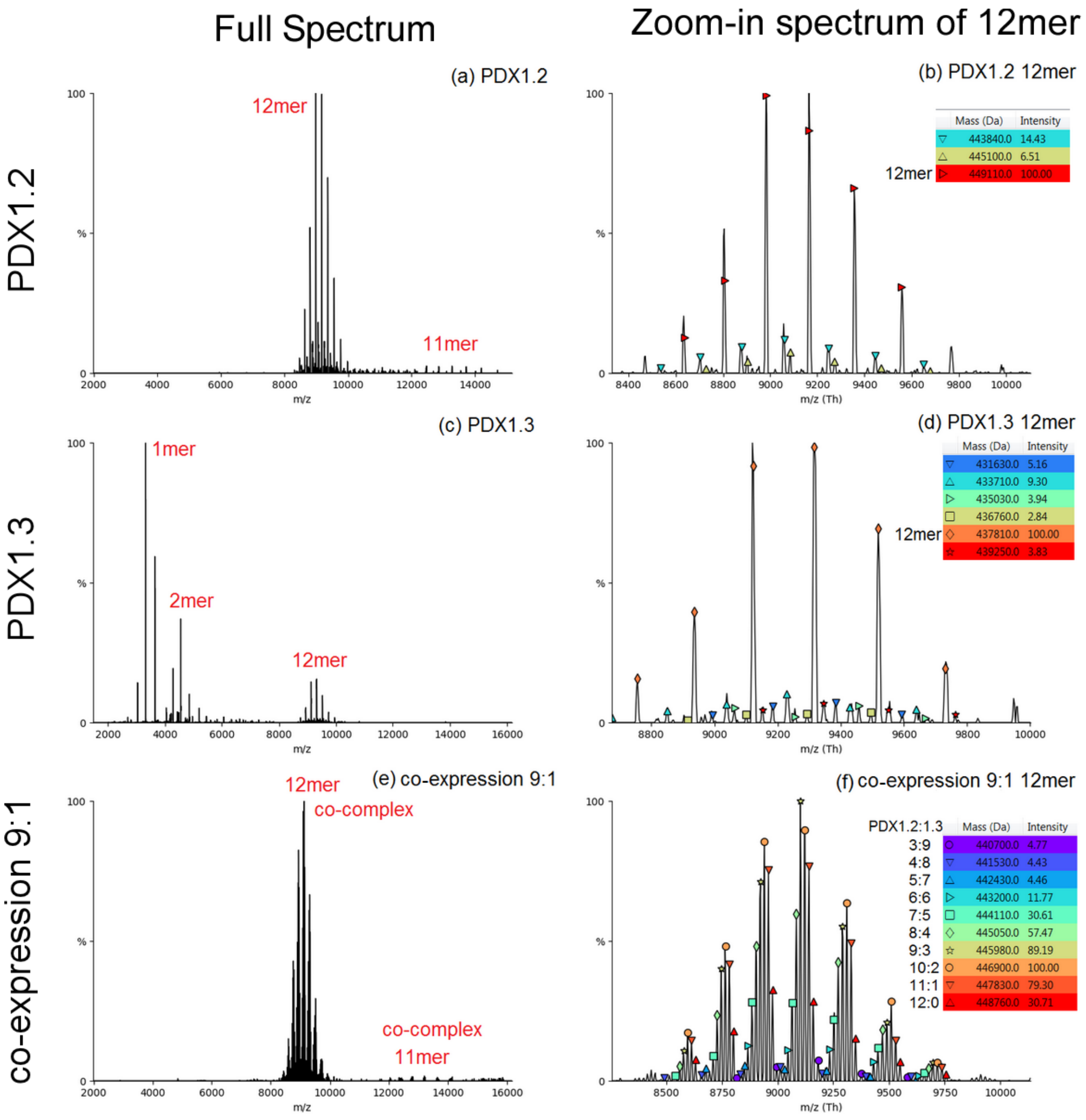

Full Spectrum
Zoom-in spectrum of 12mer
PDX1.2
PDX1.3
co-expression 9:1
(a) PDX1.2
12mer
11mer
m/z
(b) PDX1.2 12mer
Mass (Da) Intensity
443840.0 14.43
445100.0 6.51
12mer 449110.0 100.00
m/z (Th)
(c) PDX1.3
1mer
2mer
12mer
m/z
(d) PDX1.3 12mer
Mass (Da) Intensity
431630.0 5.16
433710.0 9.30
435030.0 3.94
436760.0 2.84
12mer 437810.0 100.00
439250.0 3.83
m/z (Th)
(e) co-expression 9:1
12mer co-complex
co-complex 11mer
m/z
(f) co-expression 9:1 12mer
PDX1.2:1.3 Mass (Da) Intensity
3:9 440700.0 4.77
4:8 441530.0 4.43
5:7 442430.0 4.46
6:6 443200.0 11.77
7:5 444110.0 30.61
8:4 445050.0 57.47
9:3 445980.0 89.19
10:2 446900.0 100.00
11:1 447830.0 79.30
12:0 448760.0 30.71
m/z (Th)

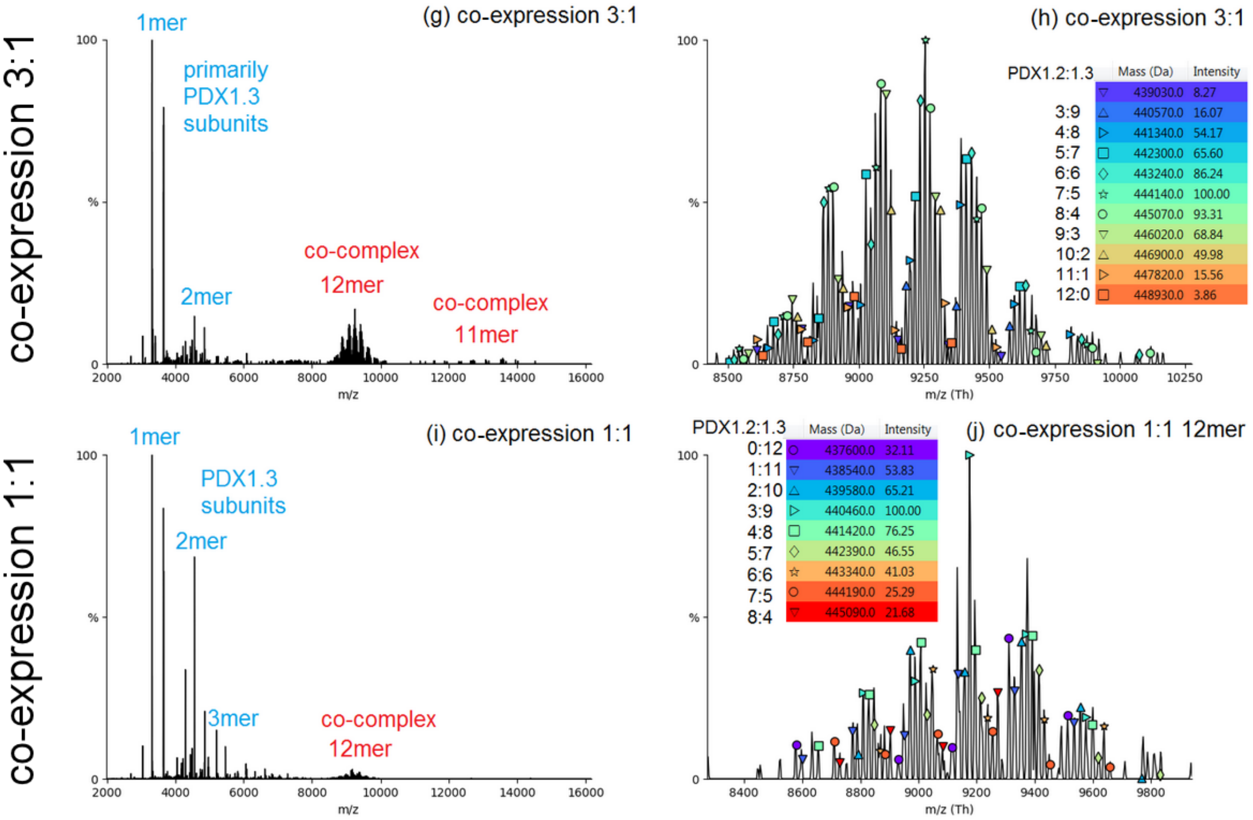

Fig. S7. (a, c, e, g, i) Full range and (b, d, f, h, j) zoomed native mass spectra of PDX1.2, PDX1.3 and hetero-complexes at co-expression conditions 9:1, 3:1 and 1:1.

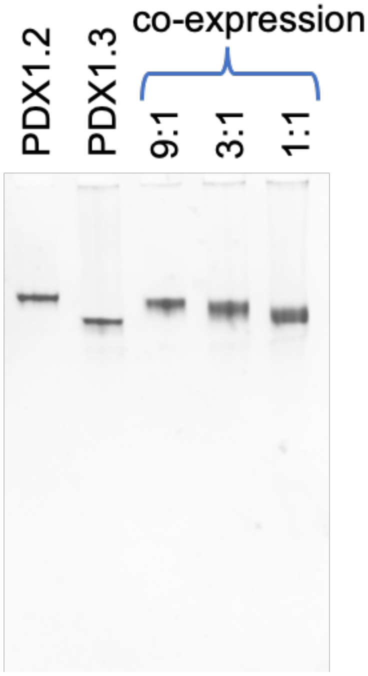


Fig. S8. Native PAGE gel of PDX complexes after buffer exchange to 200 mM ammonium acetate to verify their stability in this buffer composition. The gel was run only for 70 min on the 4-15% gradient to make sure that monomer and dimer species can be captured. No monomers or dimers are observed on the gel, suggesting that PDX1.3 homo-complex and hetero-complexes fell apart during electrospray or in the gas-phase, and not in solution.

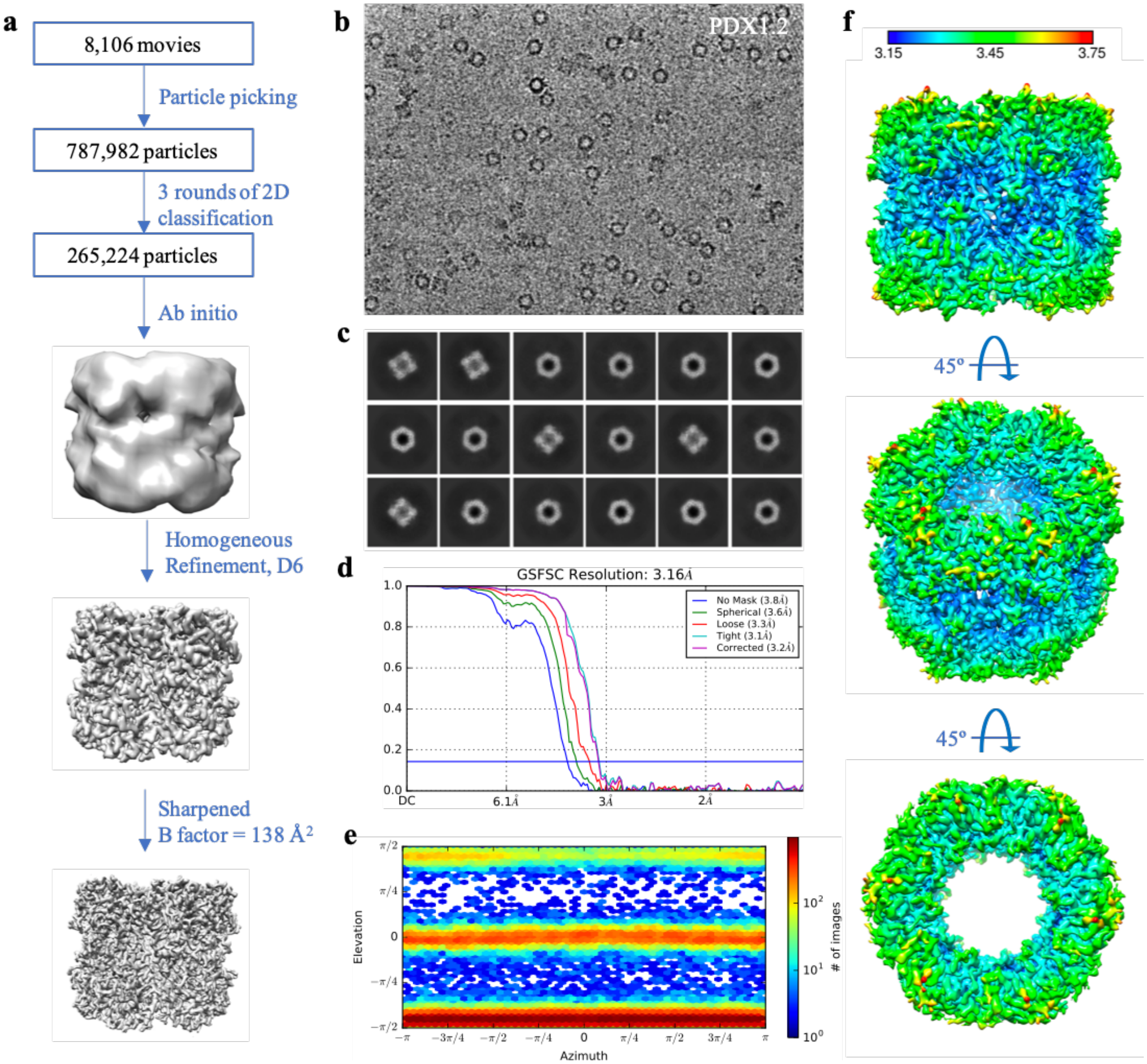


Fig. S9. Cryo-EM single particle analysis workflow for PDX1.2 homo-complex. **a**, All processing was performed in cryoSPARC v2. Movies were first corrected for beam motion, and CTF parameters were determined. The positions of particles were identified using a template-based autopicking algorithm and corrected for local motion. The final pool of 787,982 particles was subjected to three rounds of reference-free 2D classification (with 150 classes each round). The curated pool of 265,224 particles was then used for *ab initio* model generation, which was further used for homogenous refinement. The final structure shows a 3.2 Å resolution. **b**, A representative image of the PDX1.2 complexes in vitreous ice. **c**, Representative 2D classes. **d**, The Fourier shell correlation (FSC) curve and **(e)** Euler angle distribution of refined map. **f**, Sharpened PDX1.2 reconstructed map colored according to local resolution in Angstroms.

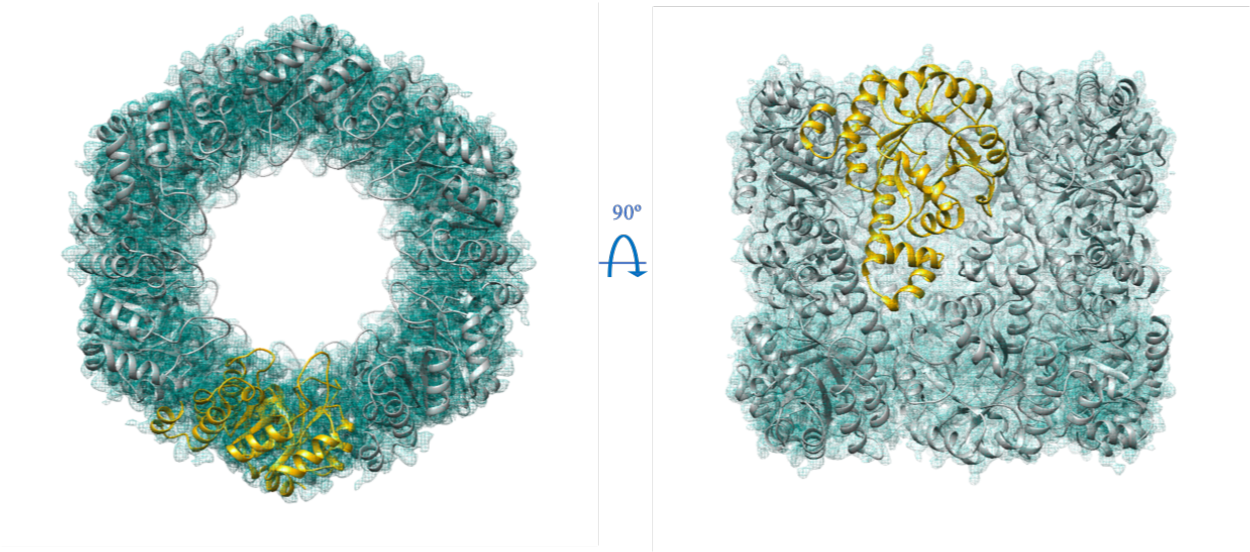


Fig. S10. Atomic model of PDX1.2 dodecamer fitted in the sharpened cryo-EM map (green mesh). An asymmetric unit (representing a single PDX1.2 subunit) is shown in yellow.

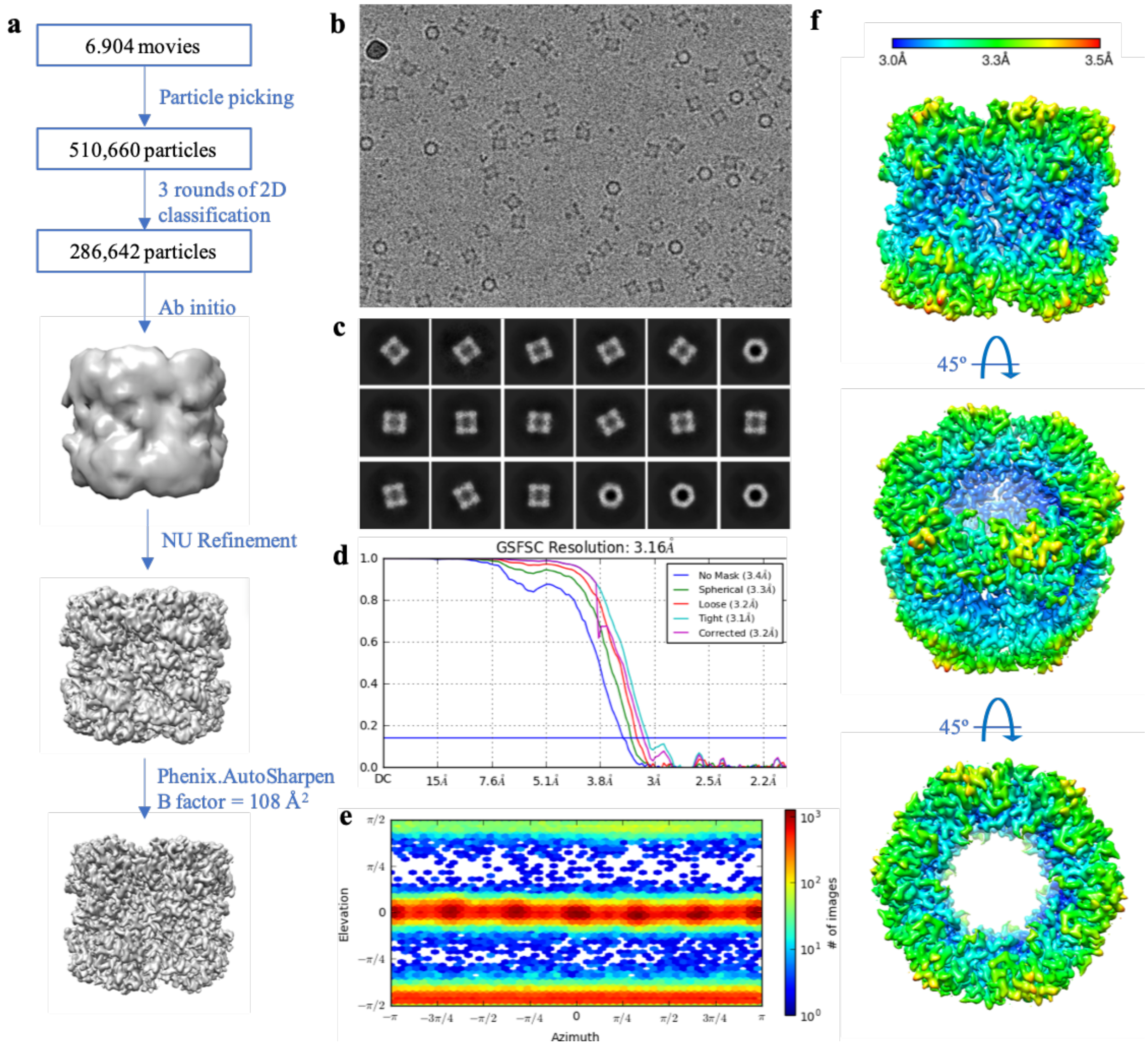


Fig. S11. Cryo-EM single particle analysis workflow for the 9:1 co-expression complex. **a**, All processing (except the sharpening) was performed in cryoSPARC v2. Movies were first corrected for beam motion, and CTF parameters were determined. The positions of particles were identified using a template-based autopicking algorithm and corrected for local motion. The final pool of 510,660 particles was subjected to three rounds of reference-free 2D classification (with 150 classes each round). The curated pool of 286,642 particles was then used in the ab initio procedure to generate an initial model, which was used for global Non-Uniform alignment. The refined structure was resolved to a 3.2 Å resolution. The sharpening of the final volume was done in PHENIX. **b**, A representative image of 9:1 co-expression complexes in vitreous ice. **c**, Representative 2D classes. **d**, The Fourier shell correlation (FSC) curve and **(e)** Euler angle distribution of refined map. **f**, Sharpened 9:1 co-expression complex reconstructed map colored according to local resolution in Angstroms

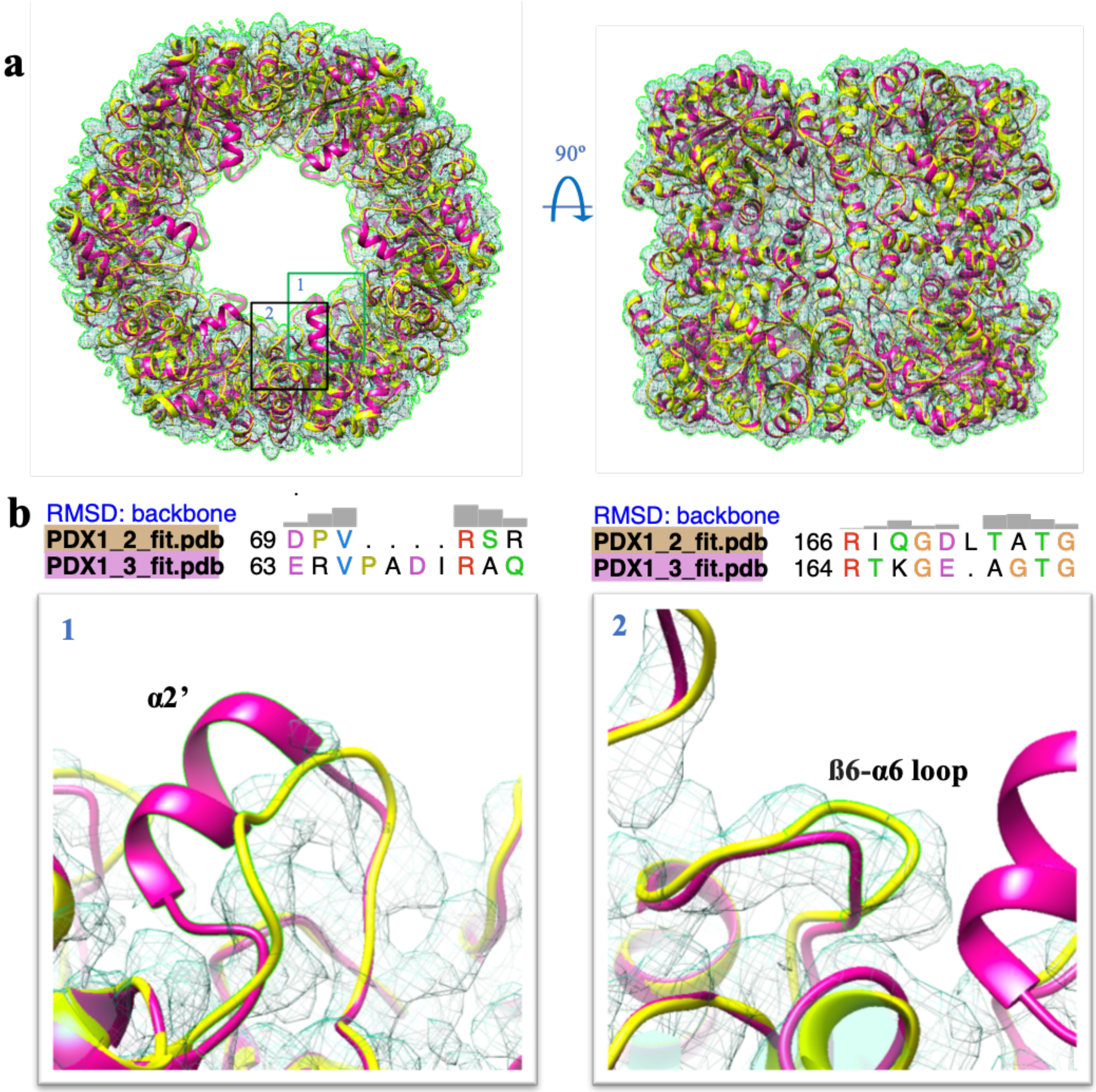


Fig. S12. Atomic modeling of PDX1.2 and PDX1.3 amino acid sequences into the cryo-EM 3D map for the 9:1 co-expression complex. **a**, Overall dodecameric fits of PDX1.2 (yellow) and PDX1.3 (magenta) proteins in the heterocomplex cryo-EM map (green mesh). Within each subunit, PDX1.2 and PDX1.3 should differ the most in the boxed areas 1 and 2 due to insertion/deletions between the two homologs. **b**, Box 1 shows the region, spanning the α2' helix for PDX1.3. Box 2 is zoomed-in area on ß6-α6 loop. Major structural differences across two proteins are defined by presence/absence of α2' helix and a ß6-α6 loop kink.

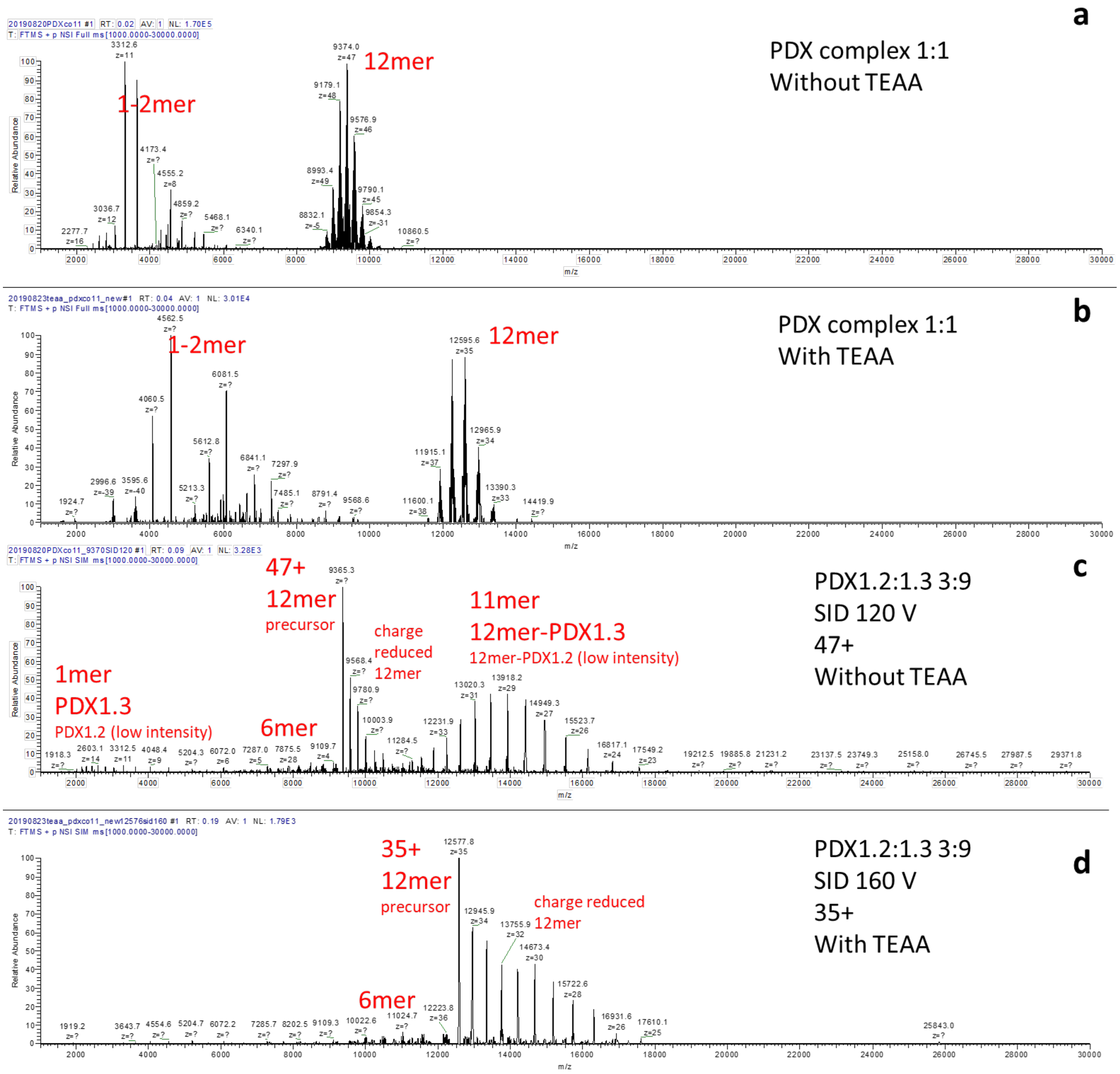


Fig. S13. Native MS of the 1:1 co-expression complex (a) without and (b) with the solution additive of triethylammonium acetate (TEAA). TEAA reduced the charge of all detected species. The most abundant charge state of the 12mer was shifted from 47+ to 35+. SID spectra of a select heteromer with the stoichiometry of 3:9 (PDX1.2:PDX1.3) (c) without and (d) with TEAA. The 1mer+11mer products were prevalent for 12mer precursor with high charge (47+), but they were not detected for lower charged precursor (35+) after TEAA addition. The dissociation pathway for generating 1mer+11mer has been associated with gas phase unfolding. Reducing the charge has been shown to suppress gas phase unfolding and allow SID to yield more structural informative products such as the 6mers *(Zhou, M., Dagan, S. & Wysocki, V. H. Impact of charge state on gas-phase behaviors of noncovalent protein complexes in collision induced and surface induced dissociation. Analyst 138, 1353-1362, doi:10.1039/c2an36525a (2013))*. Therefore, all SID experiments reported in this study were performed with reduced charge by adding TEAA, following the protocol described in the Methods section.

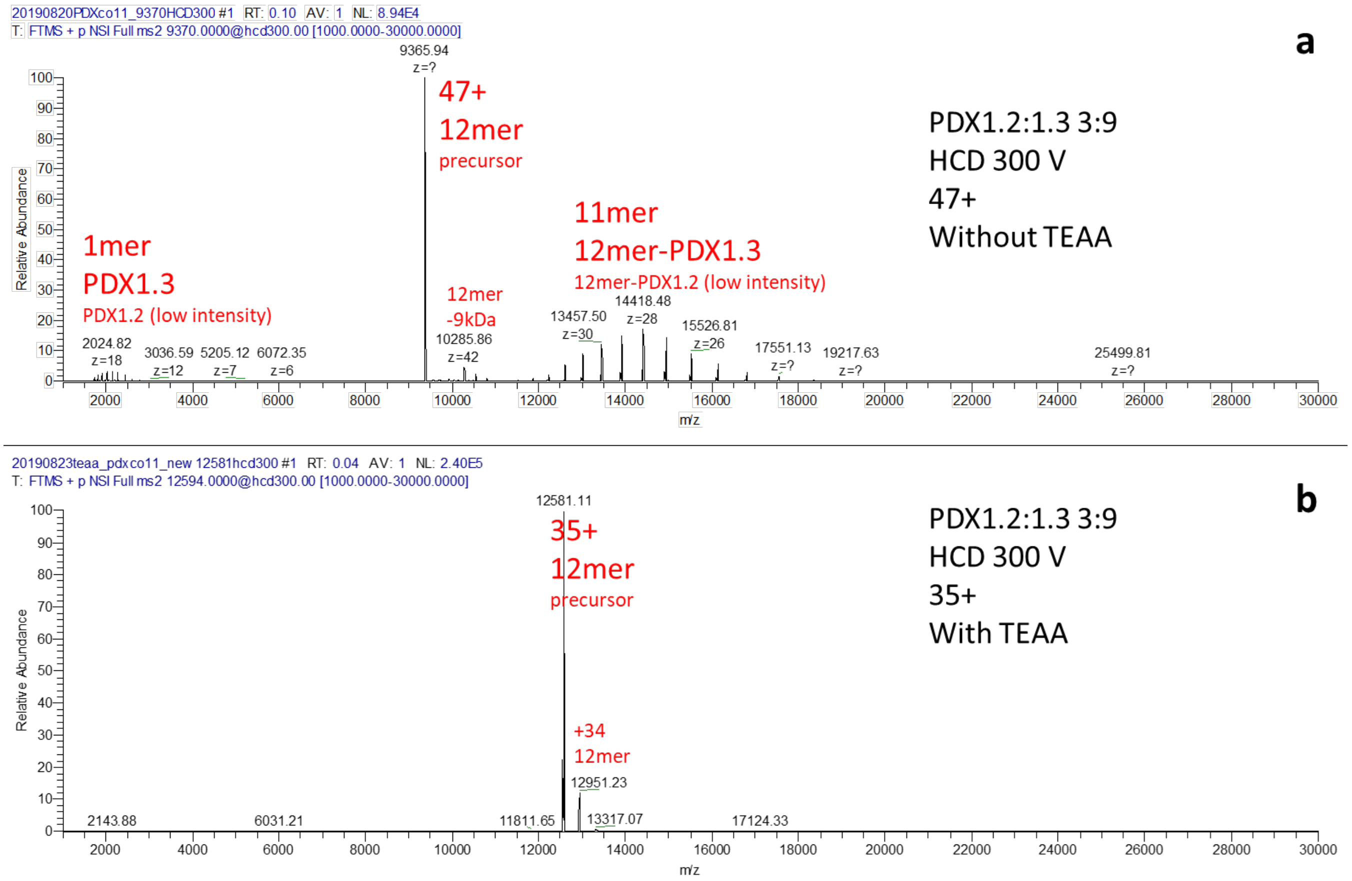


Fig. S14. HCD spectra of a select 12mer with the stoichiometry of 3:9 (PDX1.2:PDX1.3) (a) without and (b) with TEAA. Without TEAA, the 12mer dissociated into 1mer+11mer, which is the typical 1mer stripping behavior for gas phase protein dissociation upon neutral gas collision. Such behavior is generally believed to be correlated with charge directed protein unfolding in the gas phase. With TEAA, protein unfolding can be suppressed, but dissociation was also remarkedly suppressed as shown by (b). Even at the maximum collision voltage of 300 V, HCD was not able to generate significant amounts of product for charge reduced 12mer. In contrast, SID was able to yield 6mers for the charge reduced 12mer as shown in Fig. S13.

| | Deconvolted mass distribution | Raw spectra overlaid with fitted distribution of major species (excluding 12mer precursor) |
|---|---|---|

**PDX1.2**

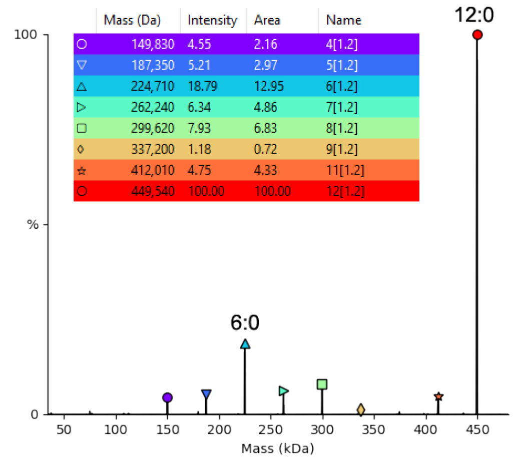


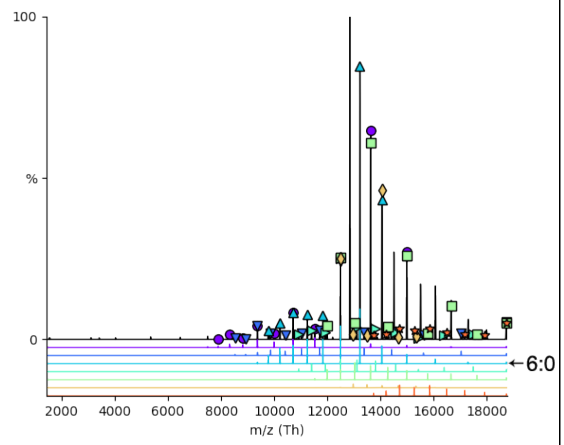


**PDX1.3**

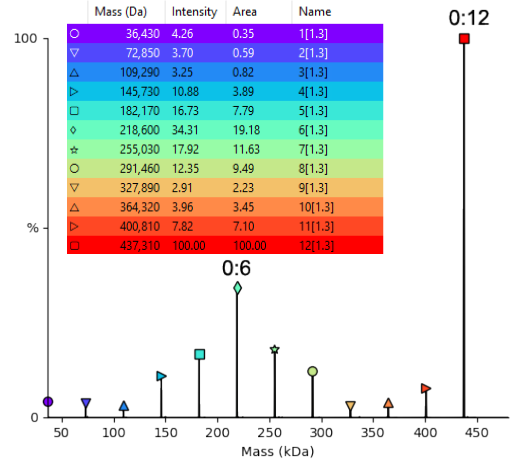


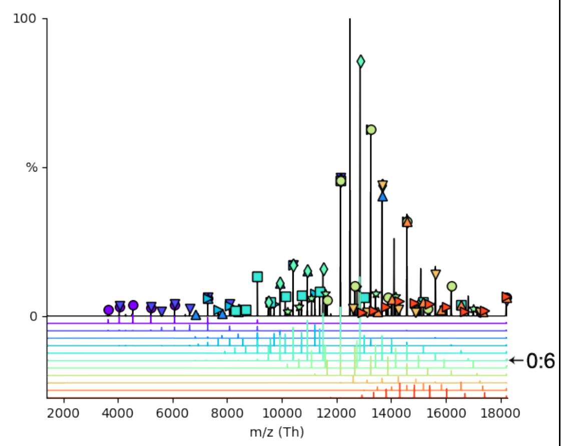


**Hetero-complex 1:11**

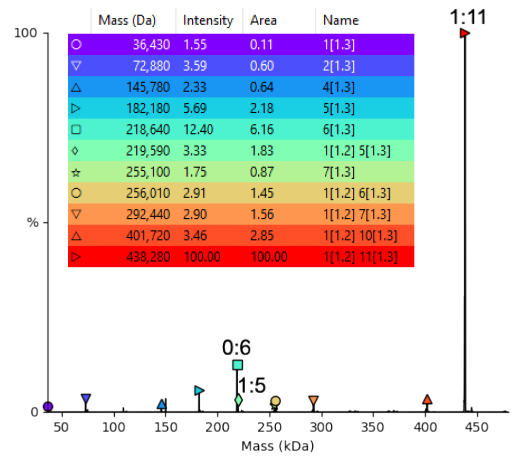


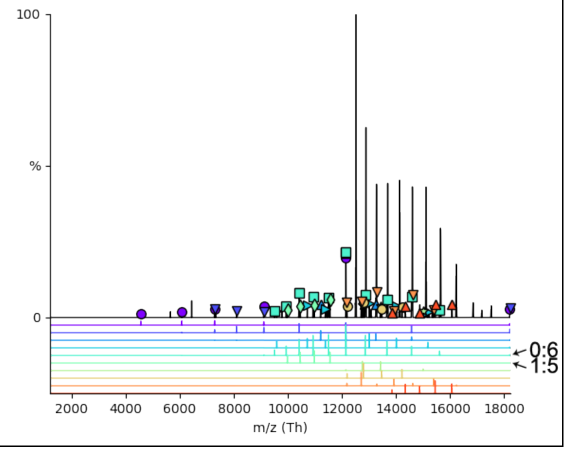

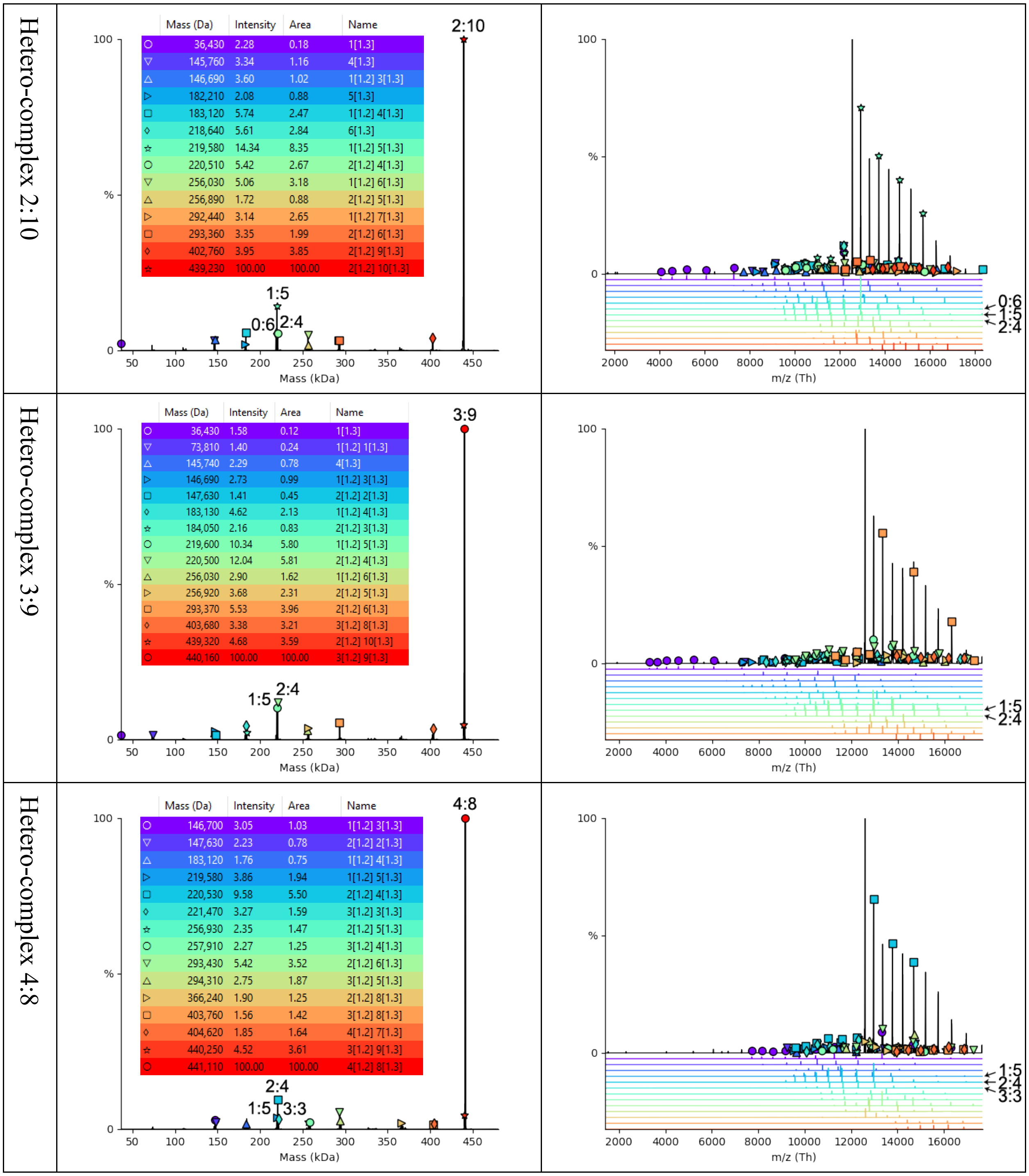
Hetero-complex 2:10
Mass (Da) | Intensity | Area | Name
36,430 | 2.28 | 0.18 | 1[1.3]
145,760 | 3.34 | 1.16 | 4[1.3]
146,690 | 3.60 | 1.02 | 1[1.2] 3[1.3]
182,210 | 2.08 | 0.88 | 5[1.3]
183,120 | 5.74 | 2.47 | 1[1.2] 4[1.3]
218,640 | 5.61 | 2.84 | 6[1.3]
219,580 | 14.34 | 8.35 | 1[1.2] 5[1.3]
220,510 | 5.42 | 2.67 | 2[1.2] 4[1.3]
256,030 | 5.06 | 3.18 | 1[1.2] 6[1.3]
256,890 | 1.72 | 0.88 | 2[1.2] 5[1.3]
292,440 | 3.14 | 2.65 | 1[1.2] 7[1.3]
293,360 | 3.35 | 1.99 | 2[1.2] 6[1.3]
402,760 | 3.95 | 3.85 | 2[1.2] 9[1.3]
439,230 | 100.00 | 100.00 | 2[1.2] 10[1.3]
2:10
1:5
0:6
2:4
Mass (kDa)
m/z (Th)
Hetero-complex 3:9
Mass (Da) | Intensity | Area | Name
36,430 | 1.58 | 0.12 | 1[1.3]
73,810 | 1.40 | 0.24 | 1[1.2] 1[1.3]
145,740 | 2.29 | 0.78 | 4[1.3]
146,690 | 2.73 | 0.99 | 1[1.2] 3[1.3]
147,630 | 1.41 | 0.45 | 2[1.2] 2[1.3]
183,130 | 4.62 | 2.13 | 1[1.2] 4[1.3]
184,050 | 2.16 | 0.83 | 2[1.2] 3[1.3]
219,600 | 10.34 | 5.80 | 1[1.2] 5[1.3]
220,500 | 12.04 | 5.81 | 2[1.2] 4[1.3]
256,030 | 2.90 | 1.62 | 1[1.2] 6[1.3]
256,920 | 3.68 | 2.31 | 2[1.2] 5[1.3]
293,370 | 5.53 | 3.96 | 2[1.2] 6[1.3]
403,680 | 3.38 | 3.21 | 3[1.2] 8[1.3]
439,320 | 4.68 | 3.59 | 2[1.2] 10[1.3]
440,160 | 100.00 | 100.00 | 3[1.2] 9[1.3]
3:9
1:5
2:4
Mass (kDa)
m/z (Th)
Hetero-complex 4:8
Mass (Da) | Intensity | Area | Name
146,700 | 3.05 | 1.03 | 1[1.2] 3[1.3]
147,630 | 2.23 | 0.78 | 2[1.2] 2[1.3]
183,120 | 1.76 | 0.75 | 1[1.2] 4[1.3]
219,580 | 3.86 | 1.94 | 1[1.2] 5[1.3]
220,530 | 9.58 | 5.50 | 2[1.2] 4[1.3]
221,470 | 3.27 | 1.59 | 3[1.2] 3[1.3]
256,930 | 2.35 | 1.47 | 2[1.2] 5[1.3]
257,910 | 2.27 | 1.25 | 3[1.2] 4[1.3]
293,430 | 5.42 | 3.52 | 2[1.2] 6[1.3]
294,310 | 2.75 | 1.87 | 3[1.2] 5[1.3]
366,240 | 1.90 | 1.25 | 2[1.2] 8[1.3]
403,760 | 1.56 | 1.42 | 3[1.2] 8[1.3]
404,620 | 1.85 | 1.64 | 4[1.2] 7[1.3]
440,250 | 4.52 | 3.61 | 3[1.2] 9[1.3]
441,110 | 100.00 | 100.00 | 4[1.2] 8[1.3]
4:8
2:4
1:5
3:3
Mass (kDa)
m/z (Th)

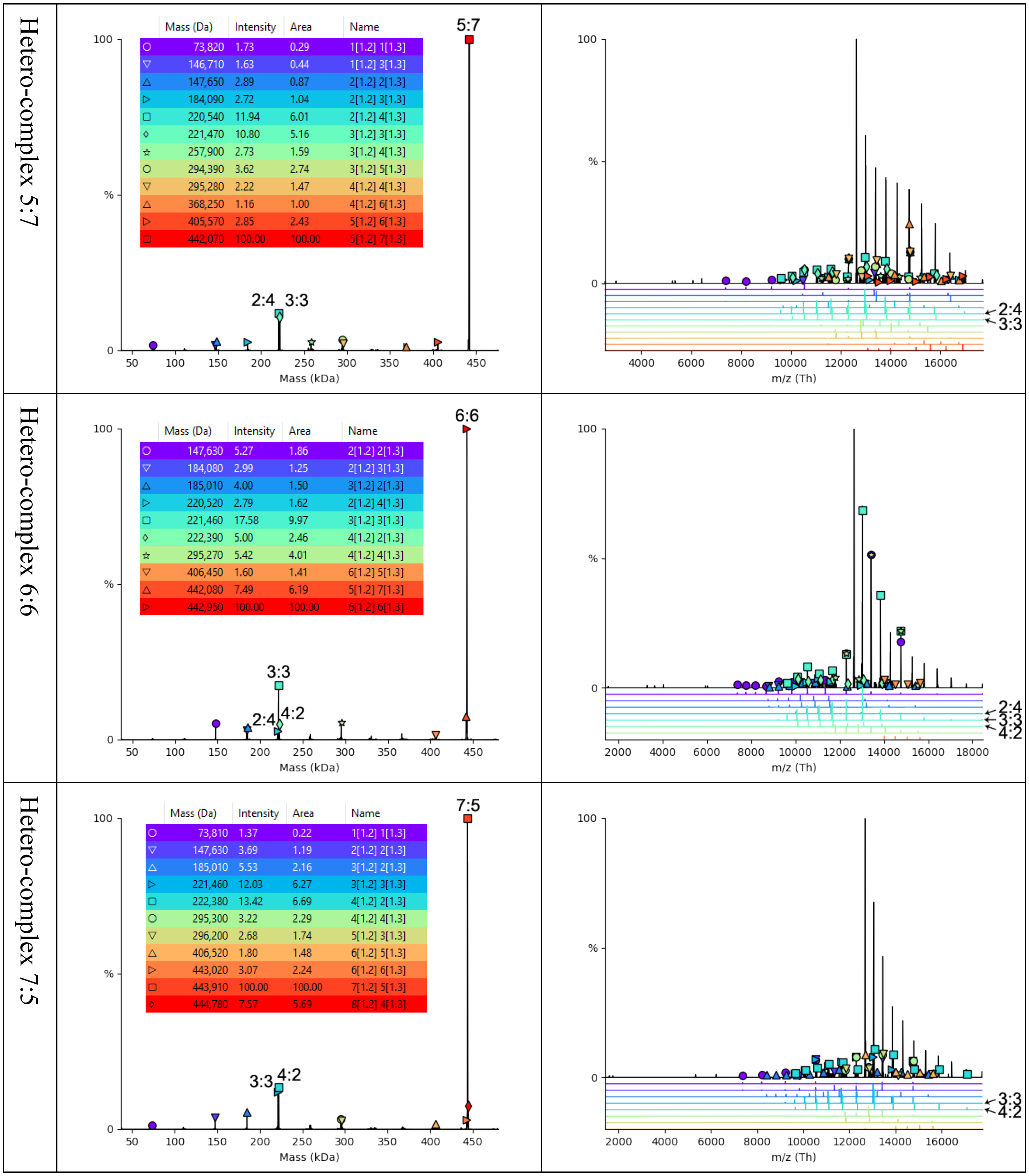
Hetero-complex 5:7
Mass (Da) Intensity Area Name
73,820 1.73 0.29 1[1.2] 1[1.3]
146,710 1.63 0.44 1[1.2] 3[1.3]
147,650 2.89 0.87 2[1.2] 2[1.3]
184,090 2.72 1.04 2[1.2] 3[1.3]
220,540 11.94 6.01 2[1.2] 4[1.3]
221,470 10.80 5.16 3[1.2] 3[1.3]
257,900 2.73 1.59 3[1.2] 4[1.3]
294,390 3.62 2.74 3[1.2] 5[1.3]
295,280 2.22 1.47 4[1.2] 4[1.3]
368,250 1.16 1.00 4[1.2] 6[1.3]
405,570 2.85 2.43 5[1.2] 6[1.3]
442,070 100.00 100.00 5[1.2] 7[1.3]
5:7
2:4 3:3
Mass (kDa)
m/z (Th)
Hetero-complex 6:6
Mass (Da) Intensity Area Name
147,630 5.27 1.86 2[1.2] 2[1.3]
184,080 2.99 1.25 2[1.2] 3[1.3]
185,010 4.00 1.50 3[1.2] 2[1.3]
220,520 2.79 1.62 2[1.2] 4[1.3]
221,460 17.58 9.97 3[1.2] 3[1.3]
222,390 5.00 2.46 4[1.2] 2[1.3]
295,270 5.42 4.01 4[1.2] 4[1.3]
406,450 1.60 1.41 6[1.2] 5[1.3]
442,080 7.49 6.19 5[1.2] 7[1.3]
442,950 100.00 100.00 6[1.2] 6[1.3]
6:6
3:3
2:4 4:2
Mass (kDa)
2:4
3:3
4:2
m/z (Th)
Hetero-complex 7:5
Mass (Da) Intensity Area Name
73,810 1.37 0.22 1[1.2] 1[1.3]
147,630 3.69 1.19 2[1.2] 2[1.3]
185,010 5.53 2.16 3[1.2] 2[1.3]
221,460 12.03 6.27 3[1.2] 3[1.3]
222,380 13.42 6.69 4[1.2] 2[1.3]
295,300 3.22 2.29 4[1.2] 4[1.3]
296,200 2.68 1.74 5[1.2] 3[1.3]
406,520 1.80 1.48 6[1.2] 5[1.3]
443,020 3.07 2.24 6[1.2] 6[1.3]
443,910 100.00 100.00 7[1.2] 5[1.3]
444,780 7.57 5.69 8[1.2] 4[1.3]
7:5
3:3 4:2
Mass (kDa)
3:3
4:2
m/z (Th)

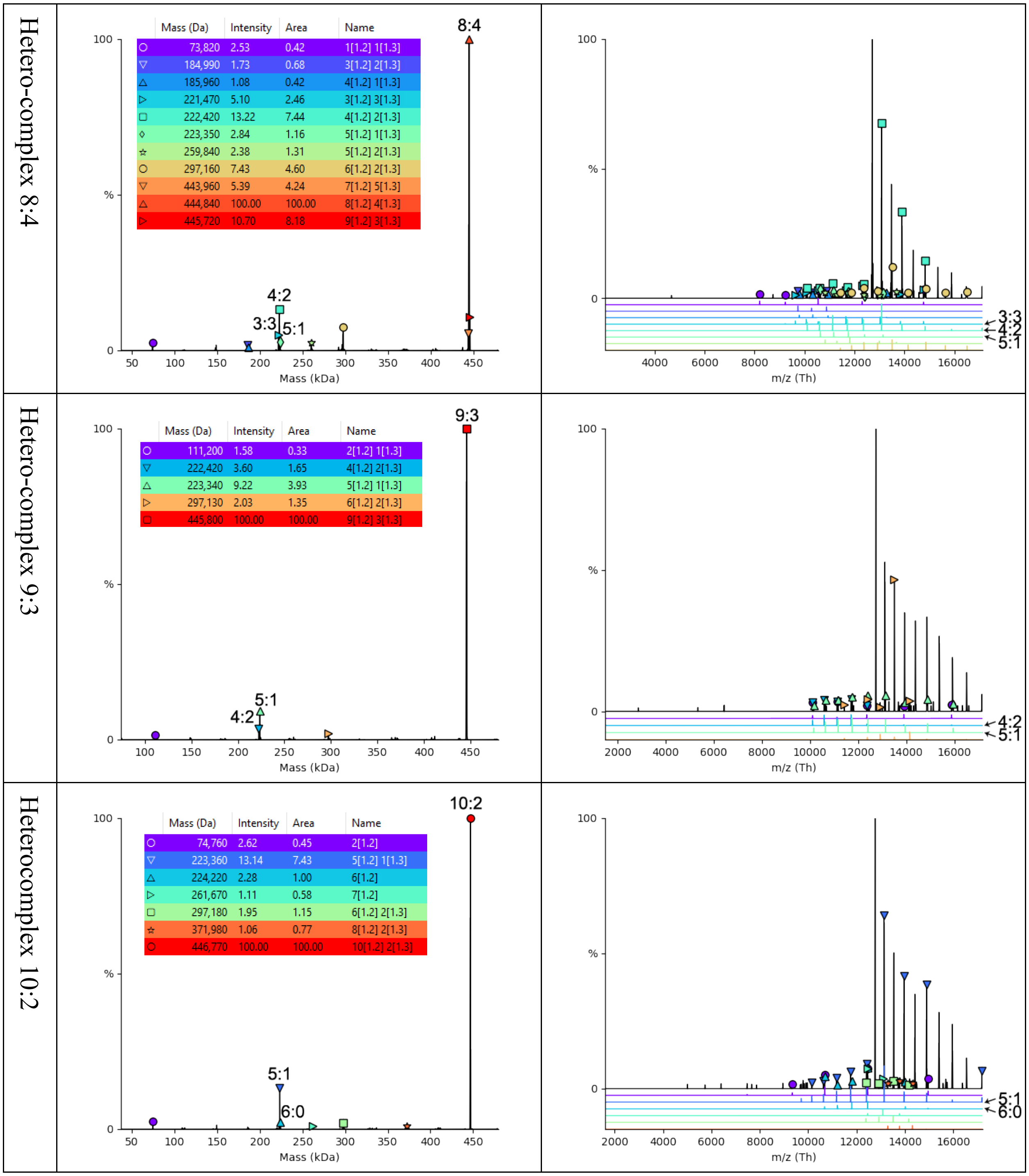
Hetero-complex 8:4
Mass (Da) | Intensity | Area | Name
73,820 | 2.53 | 0.42 | 1[1.2] 1[1.3]
184,990 | 1.73 | 0.68 | 3[1.2] 2[1.3]
185,960 | 1.08 | 0.42 | 4[1.2] 1[1.3]
221,470 | 5.10 | 2.46 | 3[1.2] 3[1.3]
222,420 | 13.22 | 7.44 | 4[1.2] 2[1.3]
223,350 | 2.84 | 1.16 | 5[1.2] 1[1.3]
259,840 | 2.38 | 1.31 | 5[1.2] 2[1.3]
297,160 | 7.43 | 4.60 | 6[1.2] 2[1.3]
443,960 | 5.39 | 4.24 | 7[1.2] 5[1.3]
444,840 | 100.00 | 100.00 | 8[1.2] 4[1.3]
445,720 | 10.70 | 8.18 | 9[1.2] 3[1.3]
8:4
4:2
3:3
5:1
Mass (kDa)
m/z (Th)
3:3
4:2
5:1
Hetero-complex 9:3
Mass (Da) | Intensity | Area | Name
111,200 | 1.58 | 0.33 | 2[1.2] 1[1.3]
222,420 | 3.60 | 1.65 | 4[1.2] 2[1.3]
223,340 | 9.22 | 3.93 | 5[1.2] 1[1.3]
297,130 | 2.03 | 1.35 | 6[1.2] 2[1.3]
445,800 | 100.00 | 100.00 | 9[1.2] 3[1.3]
9:3
5:1
4:2
Mass (kDa)
m/z (Th)
4:2
5:1
Heterocomplex 10:2
Mass (Da) | Intensity | Area | Name
74,760 | 2.62 | 0.45 | 2[1.2]
223,360 | 13.14 | 7.43 | 5[1.2] 1[1.3]
224,220 | 2.28 | 1.00 | 6[1.2]
261,670 | 1.11 | 0.58 | 7[1.2]
297,180 | 1.95 | 1.15 | 6[1.2] 2[1.3]
371,980 | 1.06 | 0.77 | 8[1.2] 2[1.3]
446,770 | 100.00 | 100.00 | 10[1.2] 2[1.3]
10:2
5:1
6:0
Mass (kDa)
m/z (Th)
5:1
6:0

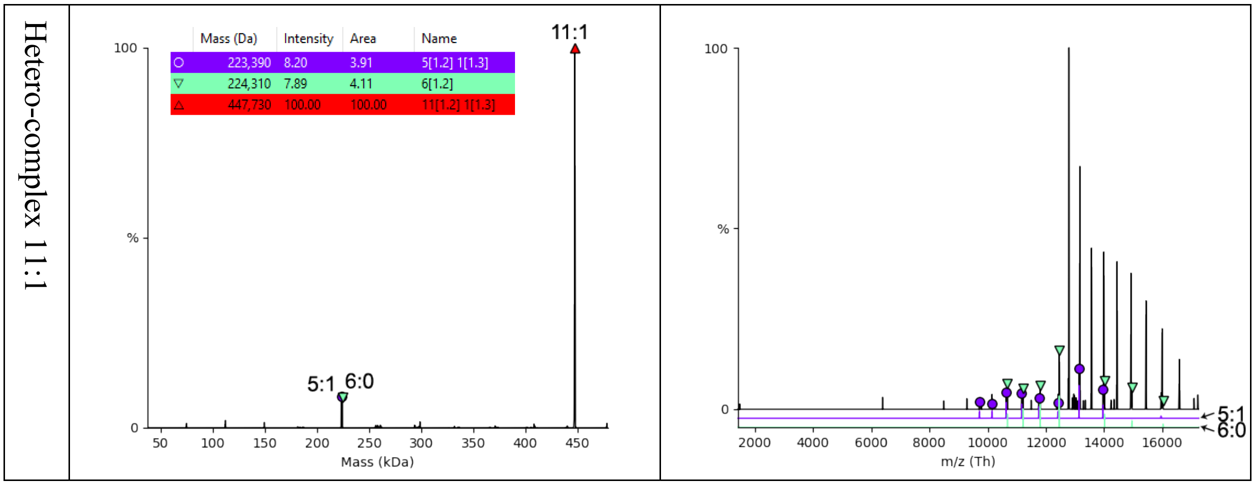


Fig. S15. Deconvoluted mass distribution (left column) and raw spectra (right column) for SID of all stoichiometry-isolated PDX 12mers (identities shown on the left) using "Oligomer and Mass Tool" in UniDec. In the deconvoluted mass distribution, major identified species are labeled with colored symbols. The keys to the symbols are shown in the inset tables, with the intensity, peak area, and assignment noted. The assignment shows the number of PDX1.2 and PDX1.3 subunits. The stoichiometry of the 12mer precursors and the 6mer products are labeled on top of the peaks (number of PDX1.2:PDX1.3). The unlabeled species at low abundance are deconvolution artifacts (bad fit with Dscore < 30% in UniDec or matched mass is lower than sequence mass). Some of the spectra contained other 12mer species at low intensity due to co-isolation. In the raw spectra (right column), peaks matching to charge state distributions are labeled with the same set of symbols. Fitted charge state distributions are also overlaid below the experimental data (black trace) and colored in the same way as the symbols. The undissociated 12mers (including charge reduced 12mers following the precursor in m/z 13000 and above with high intensity) are not labeled/shown to better display the low intensity products. The fitted charge state distributions for the 6mer species are also labeled with their assigned stoichiometry as pointed by the arrows.

**a. Theoretical: cleavage into 6mers + random assembly**

| | 0:12 | 1:11 | 2:10 | 3:9 | 4:8 | 5:7 | 6:6 | 7:5 | 8:4 | 9:3 | 10:2 | 11:1 | 12:0 |
|---|---|---|---|---|---|---|---|---|---|---|---|---|---|
| 6:0 | 0.000 | 0.000 | 0.000 | 0.000 | 0.000 | 0.000 | 0.003 | 0.020 | 0.067 | 0.222 | 0.417 | 1.000 | 1.000 |
| 5:1 | 0.000 | 0.000 | 0.000 | 0.000 | 0.000 | 0.020 | 0.090 | 0.300 | 0.533 | 1.000 | 1.000 | 1.000 | 0.000 |
| 4:2 | 0.000 | 0.000 | 0.000 | 0.000 | 0.067 | 0.300 | 0.563 | 1.000 | 1.000 | 1.000 | 0.417 | 0.000 | 0.000 |
| 3:3 | 0.000 | 0.000 | 0.000 | 0.222 | 0.533 | 1.000 | 1.000 | 1.000 | 0.533 | 0.222 | 0.000 | 0.000 | 0.000 |
| 2:4 | 0.000 | 0.000 | 0.417 | 1.000 | 1.000 | 1.000 | 0.563 | 0.300 | 0.067 | 0.000 | 0.000 | 0.000 | 0.000 |
| 1:5 | 0.000 | 1.000 | 1.000 | 1.000 | 0.533 | 0.300 | 0.090 | 0.020 | 0.000 | 0.000 | 0.000 | 0.000 | 0.000 |
| 0:6 | 1.000 | 1.000 | 0.417 | 0.222 | 0.067 | 0.020 | 0.003 | 0.000 | 0.000 | 0.000 | 0.000 | 0.000 | 0.000 |

**b. Theoretical: cleavage into 6mer rings + complete lateral symmetry**

| | 0:12 | 1:11 | 2:10 | 3:9 | 4:8 | 5:7 | 6:6 | 7:5 | 8:4 | 9:3 | 10:2 | 11:1 | 12:0 |
|---|---|---|---|---|---|---|---|---|---|---|---|---|---|
| 6:0 | 0.000 | 0.000 | 0.000 | 0.000 | 0.000 | 0.000 | 0.000 | 0.000 | 0.000 | 0.000 | 0.000 | 1.000 | 1.000 |
| 5:1 | 0.000 | 0.000 | 0.000 | 0.000 | 0.000 | 0.000 | 0.000 | 0.000 | 0.000 | 1.000 | 1.000 | 1.000 | 0.000 |
| 4:2 | 0.000 | 0.000 | 0.000 | 0.000 | 0.000 | 0.000 | 0.000 | 1.000 | 1.000 | 1.000 | 0.000 | 0.000 | 0.000 |
| 3:3 | 0.000 | 0.000 | 0.000 | 0.000 | 0.000 | 1.000 | 1.000 | 1.000 | 0.000 | 0.000 | 0.000 | 0.000 | 0.000 |
| 2:4 | 0.000 | 0.000 | 0.000 | 1.000 | 1.000 | 1.000 | 0.000 | 0.000 | 0.000 | 0.000 | 0.000 | 0.000 | 0.000 |
| 1:5 | 0.000 | 1.000 | 1.000 | 1.000 | 0.000 | 0.000 | 0.000 | 0.000 | 0.000 | 0.000 | 0.000 | 0.000 | 0.000 |
| 0:6 | 1.000 | 1.000 | 0.000 | 0.000 | 0.000 | 0.000 | 0.000 | 0.000 | 0.000 | 0.000 | 0.000 | 0.000 | 0.000 |

**c. Trimmed experiment data - 6mers moslty from major pathway**

| | 0:12 | 1:11 | 2:10 | 3:9 | 4:8 | 5:7 | 6:6 | 7:5 | 8:4 | 9:3 | 10:2 | 11:1 | 12:0 |
|---|---|---|---|---|---|---|---|---|---|---|---|---|---|
| 6:0 | 0.000 | 0.000 | 0.000 | 0.000 | 0.000 | 0.000 | 0.000 | 0.000 | 0.000 | 0.000 | 0.135 | 1.000 | 1.000 |
| 5:1 | 0.000 | 0.000 | 0.000 | 0.000 | 0.000 | 0.000 | 0.000 | 0.000 | 0.000 | 1.000 | 1.000 | 0.951 | 0.000 |
| 4:2 | 0.000 | 0.000 | 0.000 | 0.000 | 0.000 | 0.000 | 0.113 | 1.000 | 1.000 | 0.420 | 0.000 | 0.000 | 0.000 |
| 3:3 | 0.000 | 0.000 | 0.000 | 0.000 | 0.056 | 0.813 | 1.000 | 0.907 | 0.188 | 0.000 | 0.000 | 0.000 | 0.000 |
| 2:4 | 0.000 | 0.000 | 0.107 | 1.000 | 1.000 | 1.000 | 0.014 | 0.000 | 0.000 | 0.000 | 0.000 | 0.000 | 0.000 |
| 1:5 | 0.000 | 0.070 | 1.000 | 0.998 | 0.140 | 0.000 | 0.000 | 0.000 | 0.000 | 0.000 | 0.000 | 0.000 | 0.000 |
| 0:6 | 1.000 | 1.000 | 0.134 | 0.000 | 0.000 | 0.000 | 0.000 | 0.000 | 0.000 | 0.000 | 0.000 | 0.000 | 0.000 |

**d. Raw experiment data - 6mers from both major and minor pathways**

| | 0:12 | 1:11 | 2:10 | 3:9 | 4:8 | 5:7 | 6:6 | 7:5 | 8:4 | 9:3 | 10:2 | 11:1 | 12:0 |
|---|---|---|---|---|---|---|---|---|---|---|---|---|---|
| 6:0 | 0.000 | 0.000 | 0.000 | 0.000 | 0.000 | 0.000 | 0.000 | 0.000 | 0.000 | 0.000 | 0.135 | 1.000 | 1.000 |
| 5:1 | 0.000 | 0.000 | 0.000 | 0.000 | 0.000 | 0.000 | 0.000 | 0.000 | 0.156 | 1.000 | 1.000 | 0.951 | 0.000 |
| 4:2 | 0.000 | 0.000 | 0.000 | 0.000 | 0.000 | 0.000 | 0.247 | 1.000 | 1.000 | 0.420 | 0.000 | 0.000 | 0.000 |
| 3:3 | 0.000 | 0.000 | 0.000 | 0.000 | 0.289 | 0.859 | 1.000 | 0.937 | 0.331 | 0.000 | 0.000 | 0.000 | 0.000 |
| 2:4 | 0.000 | 0.000 | 0.320 | 1.000 | 1.000 | 1.000 | 0.162 | 0.000 | 0.000 | 0.000 | 0.000 | 0.000 | 0.000 |
| 1:5 | 0.000 | 0.297 | 1.000 | 0.998 | 0.353 | 0.000 | 0.000 | 0.000 | 0.000 | 0.000 | 0.000 | 0.000 | 0.000 |
| 0:6 | 1.000 | 1.000 | 0.340 | 0.127 | 0.000 | 0.000 | 0.000 | 0.000 | 0.000 | 0.000 | 0.000 | 0.000 | 0.000 |

Fig. S16. Relative abundance of the 6mer species from different 12mers based on (a) theoretical random assembly, (b) theoretical assembly with complete lateral symmetry, (c) trimmed experimental data for the 6mers from the major pathway, and (d) raw experimental data for the 6mers from all pathways. The numbers in the first columns and first rows are the PDX1.2: PDX1.3 stoichiometry of the 6mers and 12mers, respectively (similar to the format in Fig 3d). The relative abundance for (a) was calculated by counting the numbers of combinations for placing a given number (0-12) of subunits into 12 positions in a random manner. The abundances of the 6mers from the major pathway in (c) were calculated by trimming the estimated abundances of the minor pathway (described in the methods section) from raw experimental data. Relative abundance of un-trimmed data is shown in (d). Qualitatively, the raw experimental data fit between the random assembly model and the lateral symmetry model. We believe the small contribution of 6mers from the minor pathways broadened the distributions. After subtracting the contribution of the minor pathways, the “trimmed” data in (c) better fit the lateral symmetry model. Quantitatively differentiate the pathways of releasing 6mers in the experimental data require further instrument and method development. More in-depth analysis of these low abundance products will be performed in the future and will reveal additional information on the hetero-assembly.

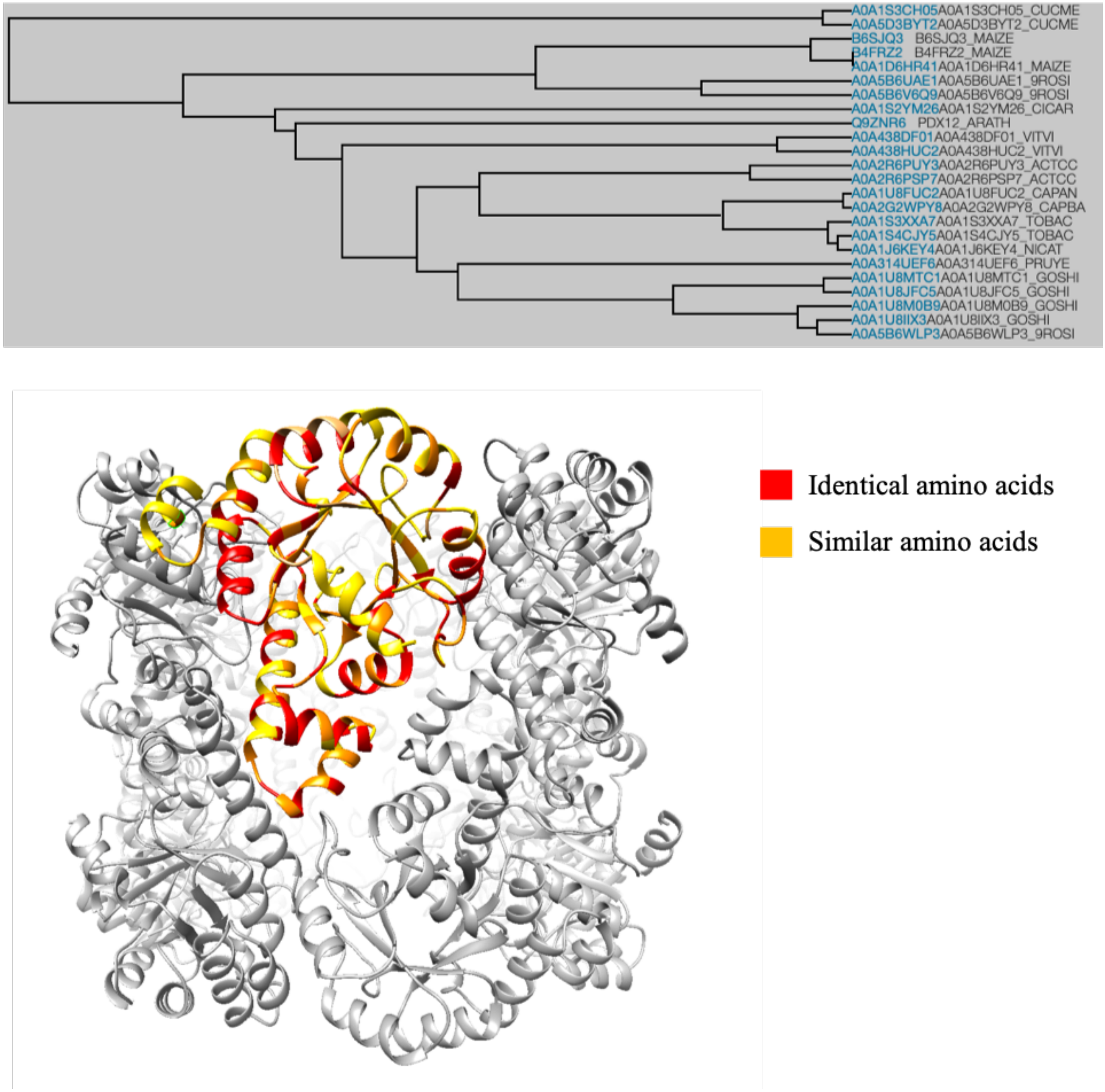


Fig. S17. Alignment of PDX1.2 sequence with pyridoxal 5-phosphate synthase-like subunit PDX1.2 sequences found in uniprot.org from other plant organisms. The individual PDX1.2 monomer was colored yellow. The most conserved residues are mapped in red and orange.

12:0

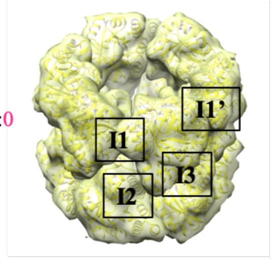


0:12

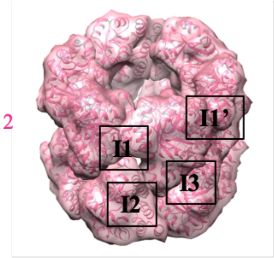


11:1

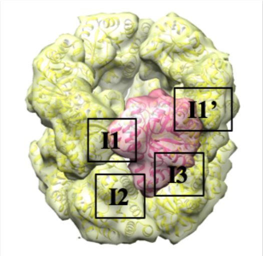


| Interface | No. of interface residues | Interface area (Å$^2$) | No. of salt bridges | No. of disulfide bonds | No. of hydrog en bonds | No. of non-bonded contacts |
|---|---|---|---|---|---|---|
| **12:0 (PDX1.2 dodecamer)** | | | | | | |
| I1=I1' (due to symmetry) | 22:18 | 931:924 | 4 | 0 | 9 | 104 |
| I2 | 7:7 | 441:443 | 4 | 0 | 6 | 37 |
| I3 | 14:14 | 909:908 | 2 | 0 | 6 | 72 |
| **0:12 (PDX1.3 dodecamer)** | | | | | | |
| I1=I1' (due to symmetry) | 17:19 | 822:851 | 3 | 0 | 2 | 58 |
| I2 | 6:6 | 420:420 | 4 | 0 | 4 | 8 |
| I3 | 12:12 | 852:851 | 3 | 0 | 2 | 52 |
| **11:1 (co-complex)** | | | | | | |
| I1 | 17:19 | 920:927 | 4 | 0 | 5 | 128 |
| I1' | 16:16 | 817:856 | 4 | 0 | 4 | 84 |
| I2 | 6:7 | 434:416 | 4 | 0 | 5 | 26 |
| I3 | 13:12 | 873:886 | 2 | 0 | 6 | 60 |

Fig. S18. Protein-protein interactions analysis by PDBsum. I1, I1', I2 and I3 denote interfaces 1, 1', 2 and 3 respectively. The hypothetical hetero-complex was constructed by structure editing via Chimera using the cryo-EM PDX1.2 atomic model and the crystal structure of PDX1.3 (PDB:5lnr).

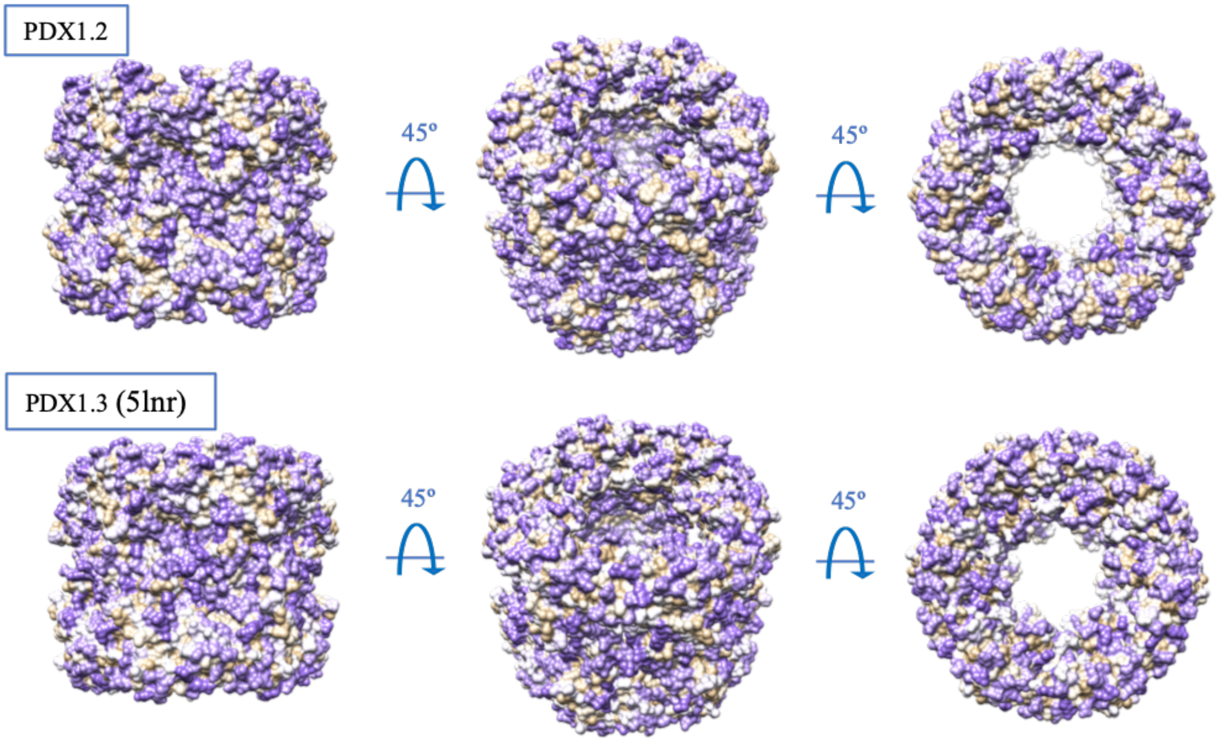


Fig. S19. Amino acid hydrophobicity plots. Purple color represents polar residues, tan color shows the most hydrophobic. Note the overall similarity in hydrophobicity while the electrostatic potential (in Fig. 4b) for these same proteins show significant differences.

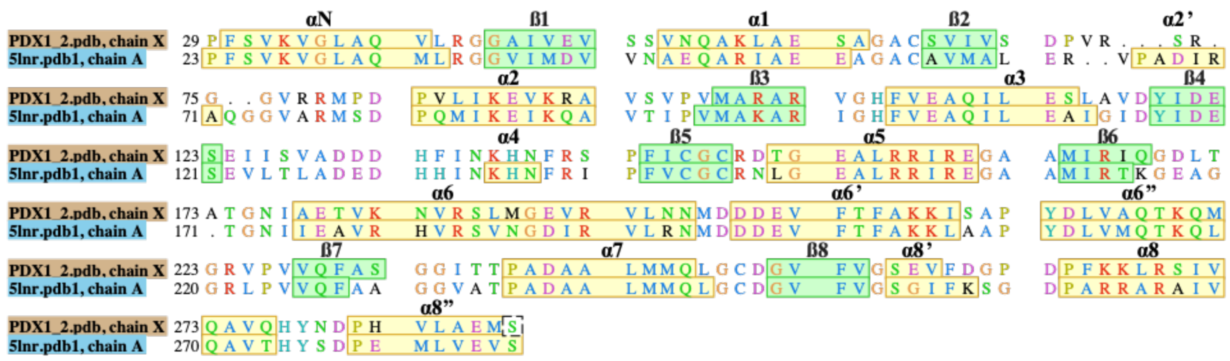


Fig. S20. Structure-based sequence alignment between PDX1.2 and PDX1.3, performed in Chimera. The residue coloring is in Clustal X default format. α-Helices (yellow) and ß-sheets (green) are annotated as herein *(Robinson, G. C. et al. Crystal structure of the pseudoenzyme PDX1.2 in complex with its cognate enzyme PDX1.3: a total eclipse. Acta Crystallogr D Struct Biol 75, 400-415, doi:10.1107/S2059798319002912 (2019)).*

| | PDX1.2 (EMD-20302, PDB 6PCJ) | 9:1 co-expression complex (EMD-20303, PDB 6PCN) | |
|---|---|---|---|
| **Data collection** | FEI Titan Krios | FEI Titan Krios | |
| Voltage (kV) | 300 | 300 | |
| Spherical aberration (mm) | 2.7 | 2.7 | |
| Total exposure dose ($e^{-}$ Å$^{-2}$) | 90 | 100 | |
| Pixel size (Å) | 0.2530 | 0.2531 | |
| **Data processing** | | | |
| Software used | cryosparc v2 | cryosparc v2 | |
| Initial number of particles | 787,982 | 510,660 | |
| Final number of particles | 265,224 | 286,642 | |
| Symmetry imposed | D6 | D6 | |
| Refinement | Homogeneous | Non-uniform (BETA) | |
| Map resolution (Å) | 3.16 | 3.16 | |
| FSC threshold | 0.143 | 0.143 | |
| 3D reference model | *ab initio* | *ab initio* | |
| **Validation** | RealSpaceRefine for PDX1.2 | RealSpaceRefine for PDX1.2 | RealSpaceRefine for PDX1.3 |
| CC (volume) | 0.82 | 0.83 | 0.84 |
| Bonds (RMSD), length (Å) | 0.003 | 0.003 | 0.004 |
| Bonds (RMSD), angles (º) | 0.495 | 0.478 | 0.585 |
| Ramachandran favored (%) | 98.06 | 97.67 | 96.81 |
| Ramachandran allowed (%) | 1.94 | 2.33 | 3.19 |
| Ramachandran outliers (%) | 0.00 | 0.00 | 0.00 |
| Rotamer outliers (%) | 0.93 | 0.00 | 0.00 |
| Cß outliers (%) | 0.78 | 0.00 | 0.00 |
| Clashscore | 7.12 | 10.19 | 10.03 |
| MolProbity score | 1.39 | 1.60 | 1.72 |

Table S1. Cryo-EM data information and PHENIX refinement statistics for PDX1.2 homomer and 9:1 co-expression complex.

| Features | PDX1.2 | 11:1 (PDX1.2: PDX1.3) | | PDX1.3 | | | |
|---|---|---|---|---|---|---|---|
| | Our model | Our model | | 5nls | | 5nlr | |
| Ligand | None | 1 PLP | apo | 12 R5P | apo | 12 PLP | apo |
| # Total amino acids | 3,120 | 3,127 | | 3,198 | | 3,198 | |
| # Arg, Lys | 348 | 349 | | 360 | | 360 | |
| # Asp, Glu | 384 | 385 | | 396 | | 396 | |
| Net charge | -36 | -36 | | -36 | | -36 | |
| Fa_elec_norm* | -2.15 | -2.15 | -2.14 | -2.19 | -2.14 | -2.15 | -2.11 |
| **Total_score_norm[▲]** | **-3.66** | **-3.67** | **-3.67** | **-3.93** | **-3.89** | **-3.87** | **-3.86** |
| Interface_delta_X[‡] | N/A (no ligand) | -11.19 (~ -134.28 if 12 PLP) | N/A (no ligand) | -236.33 | N/A (no ligand) | -155.41 | N/A (no ligand) |
| Packstat[†] | 0.66 | 0.66 | 0.66 | 0.75 | 0.74 | 0.73 | 0.73 |
| (dG_separated/dSASA) x 100[§] | -2.87 | -2.83 | -2.87 | -3.14 | -3.16 | -3.11 | -3.42 |

Table S2. Computational protein analysis by *Rosetta*. The packstat scores showed that PDX1.3 packs more tightly (0.73 ~ 0.75) than PDX1.2 (0.66). The total score also showed more favorable sum of interaction of PDX1.3 (-3.93 ~ -3.86) than PDX1.2 (~3.66). The interface stabilization (dG_separated/dSASAx100) between 12 subunits (submodels) also appears stronger in PDX1.3 (-3.42 ~ -3.11) than in PDX1.2 (-2.87). *Energy of interaction between two non-bonded charged atoms separated by distance, d. Lower values represent the more favorable energy. We present average value after normalization by the number of total amino acids. [▲] The lower the score value negatively, the protein has more stabilizing interactions. [‡] Subtracted separated energy from complex energy. The lower the interface_delta_X, the more favorable ligand binding is. [†] The higher the packstat (close to 1), the core region of protein better interdigitates leaving smaller void. [§] (Separated binding energy per unit interface area) x 100. Lower values represent the stronger complex formation from separated states.

| | Lens | Actual potential (V) |
|---|---|---|
| SID lens | Entrance 1 and 3 | -10 |
| | Entrance 2 | -2 |
| | Front top | -17.5 |
| | Front bottom | -3.5 |
| | Surface | -160 |
| | Middle bottom | -230 |
| | Back top | -274 |
| | Back bottom | -210 |
| | Exit2 | -150 |
| | Exit 1 and 3 | -170 |
| C-trap lens | Entrance lens - inject | -149 |
| | Exit lens -inject | -140 |
| | C-trap offset (upon ion injection) | -155 |

Table S3. Voltages used on the SID device in the custom modified Thermo Q-Exactive UHMR as described in previous report *(Analytical Chemistry, 91, 3611. DOI: 10.1021/acs.analchem.8b05605).*
